# Single Nuclei Sequencing Reveals Intratumoral Cellular Heterogeneity and Replication Stress in Adrenocortical Carcinoma

**DOI:** 10.1101/2024.09.30.615695

**Authors:** Liudmila V. Popova, Elizabeth A. R. Garfinkle, Daniel M. Chopyk, Jaye B. Navarro, Adithe Rivaldi, Yaoling Shu, Elena Lomonosova, John E. Phay, Barbra S. Miller, Swati Sattuwar, Mary Mullen, Elaine R. Mardis, Katherine E. Miller, Priya H. Dedhia

## Abstract

Adrenocortical carcinoma (ACC) is a rare endocrine malignancy with a poor prognosis and limited treatment options. Bulk genomic characterization of ACC has not yielded obvious therapeutic or immunotherapeutic targets, yet novel therapies are needed. We hypothesized that elucidating the intratumoral cellular heterogeneity by single nuclei RNA sequencing analyses would yield insights into potential therapeutic vulnerabilities of this disease. In addition to characterizing the immune cell and fibroblast landscape, our analyses of single nuclei gene expression profiles identified an adrenal cortex cell cluster exhibiting a program of replication stress and DNA damage response in primary and metastatic ACC. *In vitro* assessment of replication stress and DNA damage response using an ACC cell line and a series of newly-derived hormonally active patient-derived tumor organoids revealed ATR sensitivity. These findings provide novel mechanistic insight into ACC biology and suggest that an underlying dependency on ATR may be leveraged therapeutically in advanced ACC.

## INTRODUCTION

Adrenocortical carcinoma (ACC) is a rare, aggressive malignancy that originates in the adrenal cortex and has a 5-year survival of ∼6% for patients with stage IV disease.^1^ While margin negative surgical resection is the most desirable frontline treatment, many patients present with metastatic disease necessitating treatment with systemic agents. In those able to undergo resection, most experience tumor recurrence.^2–5^ Mitotane, an adrenolytic agent that also blocks cortisol synthesis, remains the only FDA-approved therapeutic agent specifically for treatment of advanced ACC. However, attaining therapeutic levels of mitotane is challenging due to significant toxicity and harmful side effects.^2,4^ Administration of chemotherapeutic drugs - etoposide, doxorubicin, and cisplatin - in conjunction with mitotane improve outcomes, but combination therapy increases median progression-free survival by only 5 months and adds significant toxicity.^2,6^ Immunotherapy has shown promise in certain cancers, yet its efficacy in ACC remains limited.^7^ Thus, there is a critical need to understand molecular mechanisms driving ACC tumor progression to improve therapy.

Bulk sequencing strategies have allowed for comprehensive genomic characterization of ACC. Previous studies using whole exome sequencing, bulk mRNA sequencing, micro RNA sequencing, DNA copy number analysis, and DNA methylation analysis have enabled molecular classifications of ACC that correlate with distinct clinical outcomes.^8,9^ Despite significant advances in understanding the genomic landscape of ACC, effective treatment options are still lacking. Single-cell RNA sequencing offers a powerful, complementary technique to bulk sequencing strategies by profiling individual cells, and thus enabling analysis of intratumoral cellular heterogeneity, a key factor that contributes to poor outcomes in many other cancers.^10–13^ The rationale for this approach in ACC includes evidence of heterogeneity in nuclear localization of β-catenin in ACC^14^ and identification of ACC tumor cell subpopulations with unique metabolomic signatures (e.g. increased pentose phosphate pathway activity) using spatial metabolomics.^15^ In addition, recent work demonstrated intratumoral heterogeneity in ACC at single-nuclei resolution.^16^ These observations suggest important cellular differences in ACC and identify a prospective cell of origin; however, they have not identified new therapeutic targets. Furthermore, prior single nuclei RNA sequencing (snRNAseq) of ACC consisted primarily of early-stage ACC, which is often amenable to surgical resection. Thus, we hypothesized that elucidating the underlying cellular biology at the single cell level in advanced ACC has the potential to identify novel therapeutic vulnerabilities in this disease. In addition, prior snRNAseq of ACC demonstrated depletion of fibroblasts and immune cells compared to normal adrenal tissue but did not fully characterize these populations to identify potential therapeutic targets.^16^ Our snRNAseq analyses of advanced ACC patient specimens further define the immune landscape and discover a population of fibroblasts associated with ACC. We also identify distinct population of adrenal cortex cells in ACC primary and metastatic samples, characterized by a unique transcriptomic program of proliferation, replication stress, and DNA damage response. Incorporating an ACC cell line and our novel ACC patient-derived tumor organoids (PTOs) in functional studies, we uncover ataxia telangiectasia and Rad3-related protein (ATR) dependency, a promising therapeutic strategy for this deadly disease.

## RESULTS

### snRNAseq reveals heterogeneous subpopulations in ACC

To profile cellular subpopulations in normal adrenal, adrenal adenoma, and ACC tissues, we performed snRNAseq on 11 specimens collected from 9 patients. The samples included normal adrenal tissue (n = 1), adrenal adenoma (AA, n = 2), primary ACC (PACC, n = 4), and metastatic ACC (MACC, n = 4, Figures 1A, 1B, and Table 1). After standard preprocessing steps, 4,735 nuclei from normal adrenal, 14,481 nuclei from AA, 23,837 nuclei from PACC samples, and 19,151 nuclei from MACC samples were used for analysis. Samples were integrated using the RPCA algorithm in Seurat and unbiased clustering resulted in 10 distinct cell populations (Figure 1C). To define adrenal cortex cells, we calculated average module scores (AC module scores) based on expression of adrenal cortex and ACC biomarkers, *NR5A1* encoding the protein SF1, *INHA*, and *MLANA*.^17–19^ This analysis identified 6 adrenal cortex clusters with high AC module scores and 4 clusters with low AC module scores. The 4 clusters with low AC module scores were identified as lymphocytes (*PTPRC*)^20^, myeloid cells (*ITGAM*)^21^, fibroblasts (*SERPINE1* and *IGFBP7*)^22,23^, and endothelial cells (*VWF* and *FLT1*).^24,25^ In addition to increased AC module scores, adrenal cortex clusters lacked expression of *PTPRC*, *ITGAM*, *SERPINE1*, *IGFBP7*, *VWF*, or *FLT1* (Figure 1D and Supplementary Figure S1). We also calculated cell type proportions and determined that normal adrenal samples were characterized by the presence of 7 different cell clusters, AA samples were characterized by the presence of 9 different cell clusters, and both PACC and MACC samples were defined by the presence of 10 different cell clusters. (Figure 1E). Overall, these data suggest that the cellular biology of ACC is defined by unique clusters when compared to normal adrenal or adrenal adenoma.

**Figure 1.**
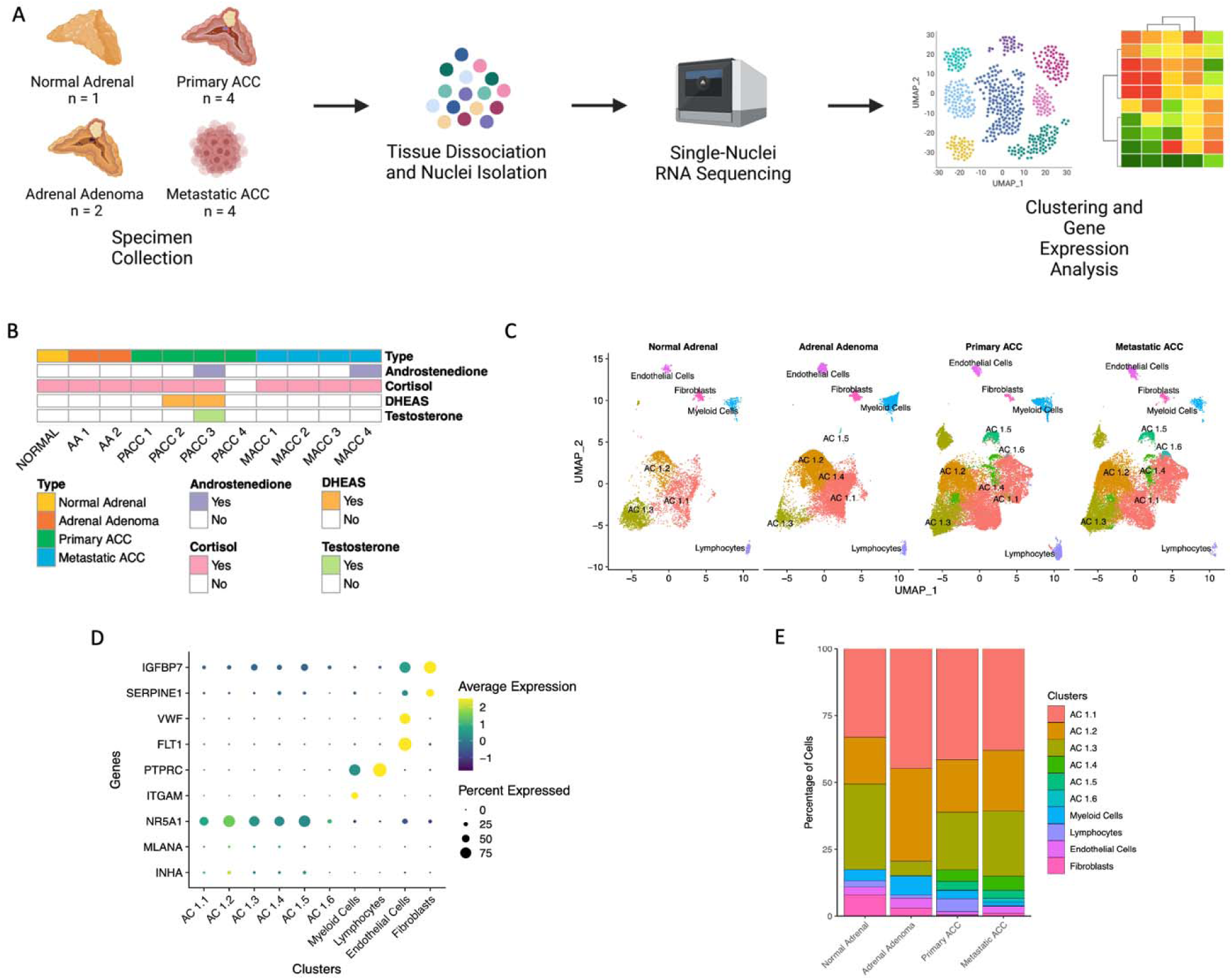
Single-Nuclei RNA Sequencing Reveals Intratumoral Heterogeneity in Normal Adrenal, Adrenal Adenoma, Primary ACC, and Metastatic ACC. A. Overview of the experimental workflow. B. Summary of the samples and their hormone secretion profiles. C. UMAP projection of the integrated Seurat object showing that 10 clusters of 5 distinct cell types were identified. D. Dot plot demonstrating marker genes that were used to annotate the identified clusters. For the dot plot, scaled average expression is displayed. E. Bar plot showing cell type proportions of the identified cell populations across different sample types. DHEAS - dehydroepiandrosterone sulfate, AC – adrenal cortex.

**Table 1.** Summary of Patient Specimens. AA – adrenal adenoma, PACC – primary ACC, MACC – metastatic ACC, DHEAS - dehydroepiandrosterone sulfate. ¹ - Foundation Medicine Testing, ² - Tempus Testing.

| Sample ID | Sample Name | Sample Description | Paired | Organ Site | Age at Surgery | Sex | Excess hormone secretion | Neoadjuvant Treatment | ENSAT Stage | Ki67 Index | Somatic Mutations | Germline Mutations |
| --- | --- | --- | --- | --- | --- | --- | --- | --- | --- | --- | --- | --- |
| Patient 1 | NORMAL | Normal Adrenal | Yes | Adrenal Gland | 35 | F | None | N/A | N/A | N/A | N/A | N/A |
|  | AA 1 | Adrenal Adenoma | Yes | Adrenal Gland |  |  | Cortisol | N/A | N/A | N/A | N/A | N/A |
| Patient 2 | AA 2 | Adrenal Adenoma | No | Adrenal Gland | 34 | M | Cortisol | N/A | N/A | N/A | N/A | N/A |
| Patient 3 | PACC 1 | Primary ACC | Yes | Adrenal Gland | 61 | M | Cortisol | Etoposide, cisplatin | 4 | No data available | No data available | No data available |
|  | MACC 1 | Metastatic ACC | Yes | Liver |  |  |  |  |  | 50% | *CHEK2* p.Ile157Thr<br>*CTNNB1* p.Ser33Cys<br>*TP53* p.Arg267Trp<br>*DUSP16-ASXL1* fusion (D3;A11*)<sup>1</sup> | No data available |
| Patient 4 | PACC 2 | Primary ACC | No | Adrenal Gland | 54 | F | Cortisol, DHEAS | Mitotane started 12 days prior to surgery | 4 | 15-20% | *DAXX* p.Glu583fs*19<sup>1</sup><br>*TP53* p.Tyr220fs*27<br>*BCORL1* p.Trp14* | No data available |
| Patient 5 | PACC 3 | Primary ACC | No | Adrenal Gland | 25 | F | Cortisol, androstenedione, testosterone, DHEAS | None | 4 | 20% | *TERT* promoter c.124C>T<sup>2</sup><br>*CDKN2A* Copy Number Loss<br>*CDKN2B* Copy Number Loss | No data available |
| Patient 6 | PACC 4 | Primary ACC | No | Adrenal Gland | 54 | F | None | None | 4 | No data available | *CTNNB1* p.Ser545Pro<br>*TP53* p.Asn200fs LOF<sup>2</sup> | No data available |
| Patient 7 | MACC 2 | Metastatic ACC | No | Liver | 57 | M | Cortisol | Mitotane | 4 | 20% (for primary tumor) | No data available | No clinically significant variants detected |
| Patient 8 | MACC 3 | Metastatic ACC | No | Lung | 45 | F | Cortisol | Mitotane | 4 | 35% (for primary tumor) | No data available | No clinically significant variants detected |
| Patient 9 | MACC 4 | Metastatic ACC | No | Cervical Lymph Node | 38 | M | Cortisol, Androstenedione | 2 months of mitotane treatment 2 years prior to surgery | 4 | 1-10% (for primary tumor) | No data available | Variants of unknown significance detected<br>*DICER* p.Trp32Cys<br>*EGLN1* p.Pro378Ser |

### ACC exhibits a unique tumor immune microenvironment

To characterize the immune landscape in adrenal tissues, we subset the lymphocyte and myeloid cell clusters (Figure 1), reprocessed, integrated using Harmony^26^, and performed unbiased reclustering. Eight immune cell types were identified including macrophages, granulocytes, regulatory T cells, CD4+ and CD8+ T cells, NK cells, B cells, and plasma cells (Figure 2A). Across all specimens, macrophages were the most common immune cell (Figure 2B, Supplementary Tables S1A-2A). Key immunocyte-specific genes were plotted in a dot plot and used to guide immune cell typing (Figure 2C).

**Figure 2.**
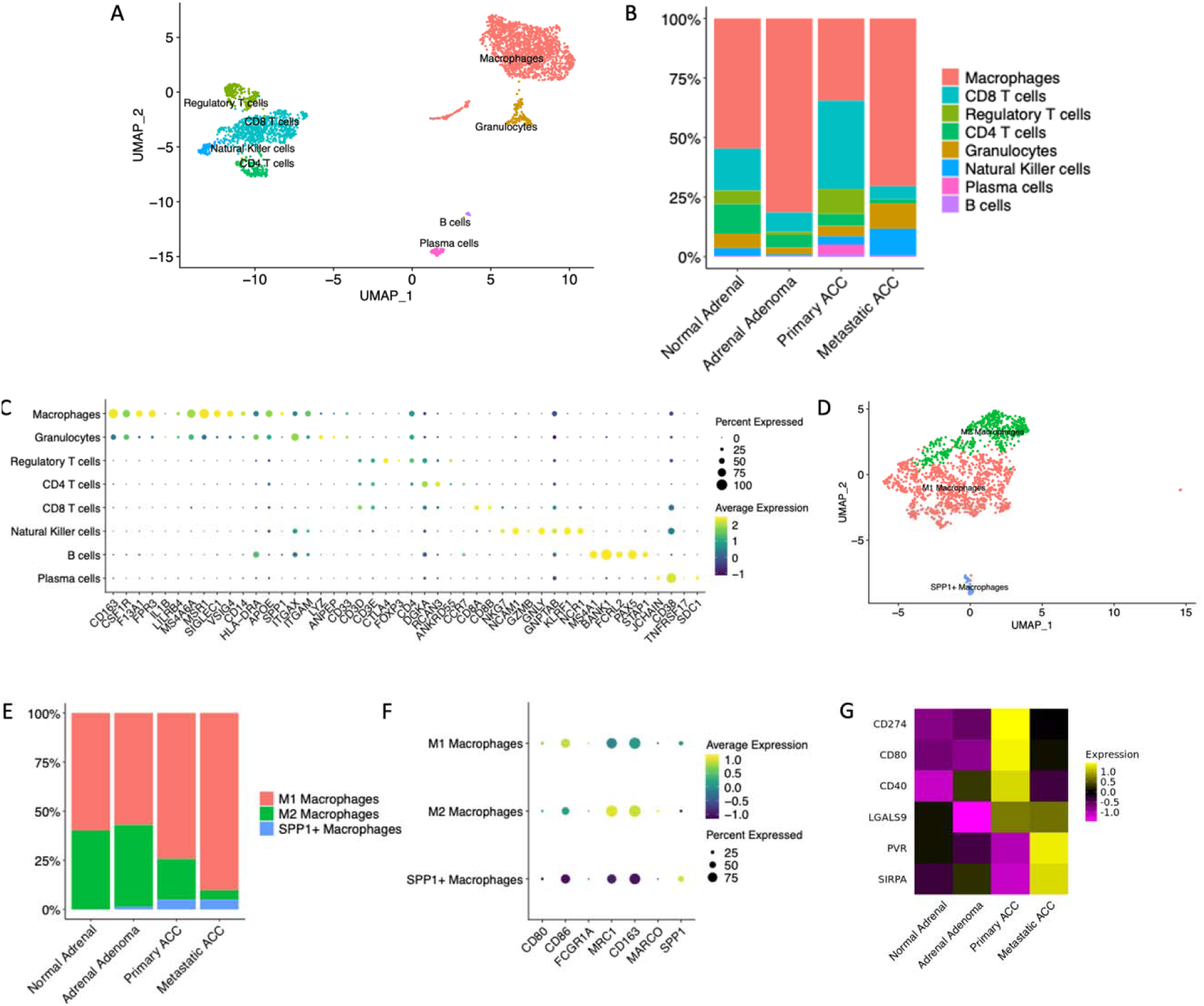
Single-Nuclei RNA-Sequencing Uncovers Infiltrating Immunocyte Populations. A. UMAP projection of subset lymphocyte and macrophage clusters from Figure 1C. B. Bar plot display of percent immunocyte composition per sample type. C. Dot plot displaying key differentially expressed cell typing markers. Scaled average expression is displayed. D. UMAP projection of subset macrophage clusters from Figure 2A. E. Bar plot display of percent macrophage subtypes per sample type. F. Dot plot displaying key differentially expressed cell typing markers. Scaled average expression is displayed. G. Heatmap displaying average expression of key macrophage immune checkpoint ligands for each sample type. Scaled average expression is displayed.

We first focused on the macrophage cluster, as these were the dominant immune population. 1,661 total macrophages were subset, reprocessed, and reclustered. From this subset, an additional 51 lymphocytes were identified and removed leaving 1,610 total macrophages for further analysis (Figure 2D). We assessed macrophage polarity by expression of proinflammatory or M1 markers, *CD80*, *CD86*, and *FCGR1A* (also known as *CD64*),^27,28^ and immunosuppressive or M2 markers, *MRC1*, *CD163*, and *MARCO*^29–31^. Although we noted no significant differences in the proportion of M1 macrophages across all tissues, we found a significant decrease in M2 macrophages in PACC and MACC specimens compared to normal and AA specimens (false discovery rate (FDR) <0.05, Figure 2E). Prior work correlated increased M2 macrophage infiltration with worse prognosis in other solid tumors; however, more recent evidence indicates plasticity in M1 and M2 polarization.^32^ Furthermore, macrophage expression of *SPP1* was strongly associated with clinical prognosis independent of classic M1 versus M2 macrophage markers.^33,34^ Additionally, SPP1+ macrophages have been associated with an immunosuppressive phenotype.^32,35,36^ Therefore, we assessed *SPP1* expression and found a significant increase in SPP1+ macrophages (FDR = 0.03) in PACC and MACC specimens compared to normal and AA (Figure 2E, Supplementary Tables S1B-2B). Our findings suggest that SPP1+ macrophages are associated with ACC tumor progression.

We next analyzed the expression of immune checkpoint molecules expressed by macrophages to identify potential strategies for immunotherapy. We plotted the average expression of pro-inflammatory (*CD40* and *CD80*) and immunosuppressive (*CD274* or *PDL1*, *LGALS9*, *PVR*, *SIRPA,* and *CD80*) markers (Figure 2G).^37–44^ CD80 has a dual role as either a costimulatory or coinhibitory ligand depending on whether it binds CD28 or CTLA-4, respectively, on neighboring T cells.^44^ We found that PACC-associated macrophages had higher average expression of *CD274,* and *LGALS9*, and MACC-associated macrophages had higher average expression of *PVR*, and *SIRPA.* PACC-associated macrophages also exhibited elevated expression of *CD40* and *CD80* relative to the benign tissues, suggesting these markers may serve as potential targets for immunotherapy. Taken together, our data suggest a changing macrophage landscape from normal adrenal to ACC that increasingly presents an immunosuppressive environment.

Analysis of lymphoid populations demonstrated that both T and NK cells were reduced or absent in most ACC specimens. Interestingly, PACC 4, a tumor from a patient without clinical or biochemical evidence of excess cortisol secretion, had the highest numbers of T and NK cells compared to other samples (Supplementary Table S2A). This finding is consistent with the association between hypercortisolism and reduced T cell infiltration in ACC (Table 1 and Figure 1B).^8,45^ Therefore, our analysis of CD8+, CD4+, and NK cells were primarily represented by PACC 4. First, we subset CD8+ T cells and plotted average expression of key immunoregulatory molecules (Supplementary Figure S2). Our analysis revealed that CD8+ T cells in the ACC samples compared to those in benign tissue had reduced expression of markers associated with immune activating functions (*CD28* and *CD40L*G) and increased expression of markers of T cell exhaustion (*CTLA-4*, *PDCD1*, *TIGIT*, *LAG3*, and *HAVCR2*).^46–48^ We next assessed expression of these markers within the CD4+ T cell subset and found that expression of *CD28*, *CTLA-4*, *TIGIT*, and *LAG3* was increased whereas expression of *CD40LG*, *PDCD1*, and *HAVCR2* was decreased in ACC relative to normal adrenal and AA (Supplementary Figure S2). Taken together, our analyses demonstrate signs of T cell exhaustion in ACC which was more pronounced in CD8+ compared to CD4+ T cells. Finally, analysis of the NK cells revealed increased expression of *TIGIT*^49^ and the NK inhibitory receptor *KLRC1*^50^ in ACC relative to normal adrenal and AA specimens (Supplementary Figure S2). Collectively, these findings indicate that hypercortisolism in ACC reduces T cell infiltration, and thus ACC is characterized by an immunosuppressive microenvironment, which may require precision therapeutic strategies that enhance immune infiltration in combination with immunotherapy to achieve efficacy.

### ACC is associated with a unique fibroblast subpopulation

We next subset fibroblasts to determine if malignant specimens are associated with changes in fibroblast subpopulations. Initially we identified 10 fibroblast clusters. Four clusters including adipocytes were excluded from further analysis, and the remaining fibroblast data were reprocessed and reintegrated to identify 5 fibroblast clusters (Figure 3A-B). These clusters were annotated based on marker gene expression (Figure 3C) and fibroblast markers (Figure 3D).^51,52^ Fibroblast cluster 3.1 was present in all specimens and was the predominant cluster in PACC and MACC (Figure 3B). Characterized by high expression of *ACTA2*, *MMP11*, and *HAS2,* cluster 3.1 likely represents myofibroblastic cancer-associated fibroblasts (myCAFs).^53,54^ Further analysis of fibroblast cluster 3.1 in PACC specimens for known cancer-associated fibroblasts markers^55^ demonstrated increased levels of *PDGFRB, FAP, VIM, POSTN, HAS2, CXCL8, CXCL2, LMNA, and HAS 1.* MACC specimens expressed increased *CAV1*, *CFD*, *MMP11*, *COL1A1*, which are additional markers of cancer-associated fibroblasts (Figure 3E). These findings suggest that ACC is characterized by the expansion of a distinct fibroblast subpopulation at both primary and metastatic sites.

**Figure 3.**
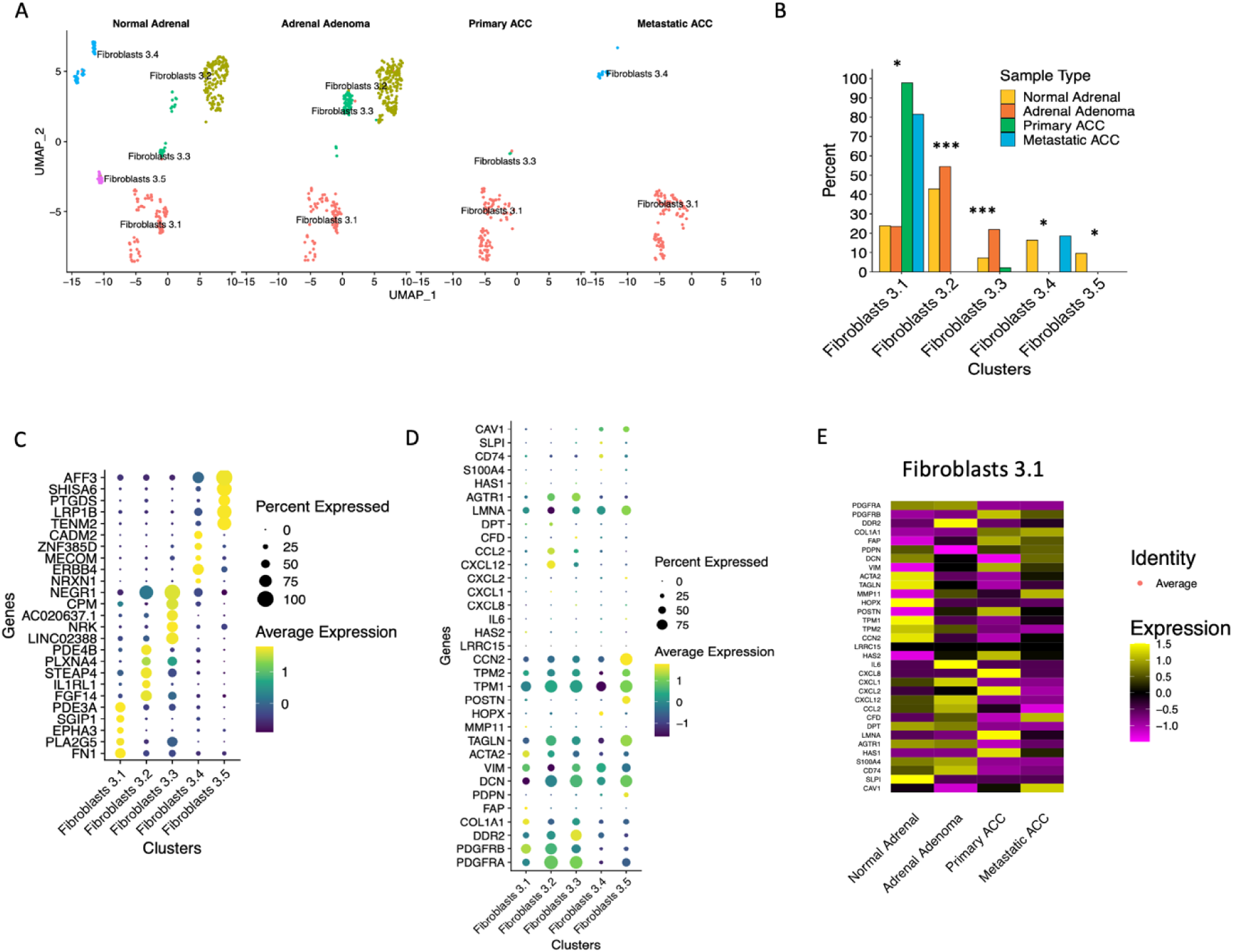
Single-Nuclei Sequencing Identifies Cancer-Associated Fibroblasts in ACC. A. UMAP projection demonstrating that further subclustering of fibroblasts identified in Figure 1C results in identification of 5 fibroblast clusters, denoted 3.1 through 3.5. B. Bar plot demonstrating cell type proportions of each of the identified fibroblast clusters ( * - FDR <= 0.05, ***-FDR <= 0.001), C. Dot plot demonstrating top marker genes which were used to annotate the identified clusters. Top 5 markers per cluster are displayed. D. Dot plot demonstrating expression of known of fibroblast marker genes. For both dot plots, scaled average expression is displayed. E. Heatmap showing expression of selected fibroblast markers in fibroblast cluster 3.1. For the heatmap, scaled average expression is displayed. FDR – false discovery rate.

### Adrenal cortex clusters demonstrate unique transcriptional programs

To further characterize the 6 adrenal cortex clusters, we subset, reintegrated, and performed unbiased clustering of these data, which resulted in identification of 7 adrenal cortex clusters and one adrenal medulla cluster (defined by *CHGA* expression, Supplementary Figure S3).^56^ Adrenal medulla cells were predominantly found in normal adrenal tissue (150 cells). After excluding adrenal medulla cells, we reintegrated and reclustered adrenal cortex cells to reveal 7 final adrenal cortex clusters (AC 3.1-3.7) based on marker gene expression (Figures 4A and B). The majority of the cells in cluster 3.7 originated from PACC 3, the only specimen with excess DHEAS, androstenedione, and testosterone secretion. Consistent with these clinical findings, AC 3.7 expressed high levels of *CYP11A1* and *CYP17A1* which are involved in testosterone, DHEAS, and androstenedione production. (Figure 4C, Table 1, and Supplementary Figure S4).^57,58^ We then assessed subpopulation frequency across different sample types and found that clusters AC 3.5 and AC 3.6 were significantly enriched in PACC and MACC specimens (FDR < 0.0001, Figure 4D). In order to identify gene expression changes associated with tumor progression, we calculated average module scores for each of the clusters for Hallmark pathways downloaded from MSigDB^59,60^ (Figure 4E and Supplementary Figure S5). In these clusters, in which cell cycle regression was performed, we found that cluster AC 3.6 was characterized by the highest module scores for the following hallmark pathways: E2F targets, G2M checkpoint, mitotic spindle, PI3K AKT mTOR signaling, and DNA repair. Hence, AC 3.6 bears a gene expression signature of dysfunctional DNA replication and repair, indicating potential mechanistic underpinnings of malignancy and tumor progression. Cluster AC 3.5 was defined by TGF-β signaling, unfolded protein response, and glycolysis. The unfolded protein response and glycolysis signatures in cluster 3.5 were enriched only in MACC, whereas the DNA damage repair signature in cluster AC 3.6 was present in all PACC and MACC samples (Supplementary Figure S5). We also found that cluster AC 3.3 was defined by the Wnt/β catenin signaling pathway, which is commonly dysregulated in ACC.^2^ Cluster AC 3.2 was defined by cholesterol homeostasis, indicating a potential role in steroid metabolism.

**Figure 4.**
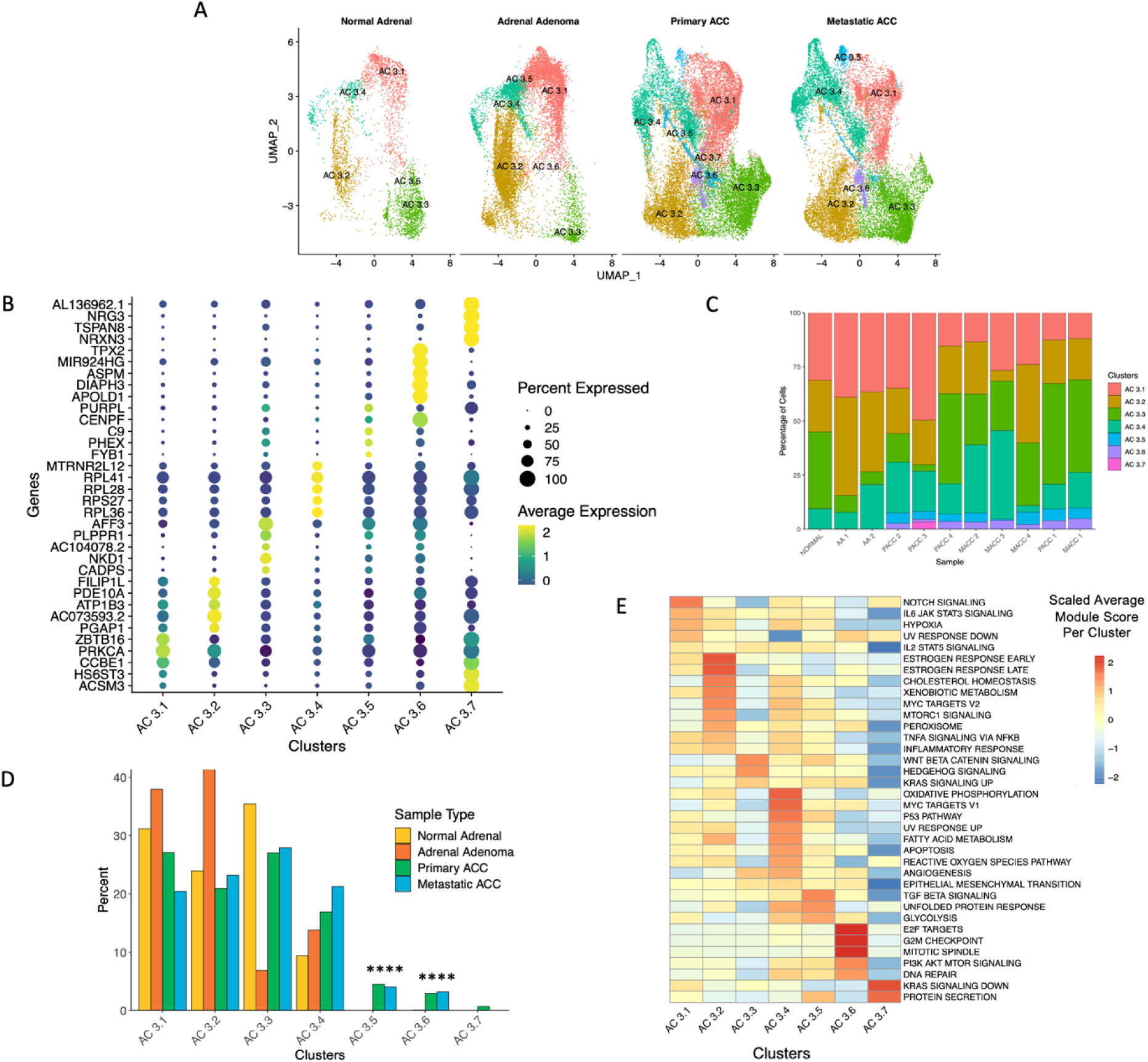
Single-Nuclei RNA Sequencing Demonstrates Intratumoral Heterogeneity of Adrenal Cortex Cells. A. UMAP projection showing that further subclustering of adrenal cortex cells results in identification of 7 adrenal cortex clusters, denoted AC 3.1 through AC 3.7. B. Dot plot demonstrating top 5 marker genes which were used to annotate each of the identified clusters. For the dot plot, scaled average expression is displayed. C. Bar plot showing cell type proportions of each of the adrenal cortex clusters (*** - FDR <= 0.001). D. Bar plot demonstrating cell type proportions of each of the identified clusters on a per-sample basis. E. Heatmap demonstrating average module scores for a selection of Hallmark pathways downloaded from MSigDB for each of the identified clusters. In the heatmap, average module scores are scaled on a per-row basis.

### Replication stress and DNA damage repair signature is enriched in cluster AC 3.6

To better understand the role of replication stress and DNA damage repair pathways in ACC, we used REACTOME pathways downloaded from the MSigDB database to generate module scores for each of the identified clusters.^59,61^ We observed that cluster AC 3.6 in PACC and MACC was characterized by the highest module scores associated with activation of ATR in response to replication stress, homology-directed repair, non-homologous end joining, mismatch repair, and base excision repair (Figure 5A). Additionally, we observed that replication stress and DNA repair pathways were enriched in PACC and MACC specimens (Supplementary Figure S6). We next used QIAGEN Ingenuity Pathway Analysis (IPA)^62^ to compare cluster AC 3.6 to other clusters found in PACC and MACC specimens and again identified that cell cycle checkpoints and homology-directed repair through homologous recombination or single-strand annealing were among the top 10 upregulated pathways (Figure 5B). Replication stress and chromosome instability have been previously associated with tumorigenesis and tumor progression.^63^ To elucidate genomic instability in AC clusters, we used inferCNV^64^ to identify copy-number variation (CNV) in tumor specimens on a per-cluster basis. We found that cluster AC 3.6 had the highest total number of CNVs, including both copy number gains and losses compared to other clusters (Figure 5C-E). This observation suggests that cluster AC 3.6, which demonstrates dysregulated replication stress and DNA damage repair response as well as genomic instability, may reflect the underlying biology driving ACC tumor development and progression.

**Figure 5.**
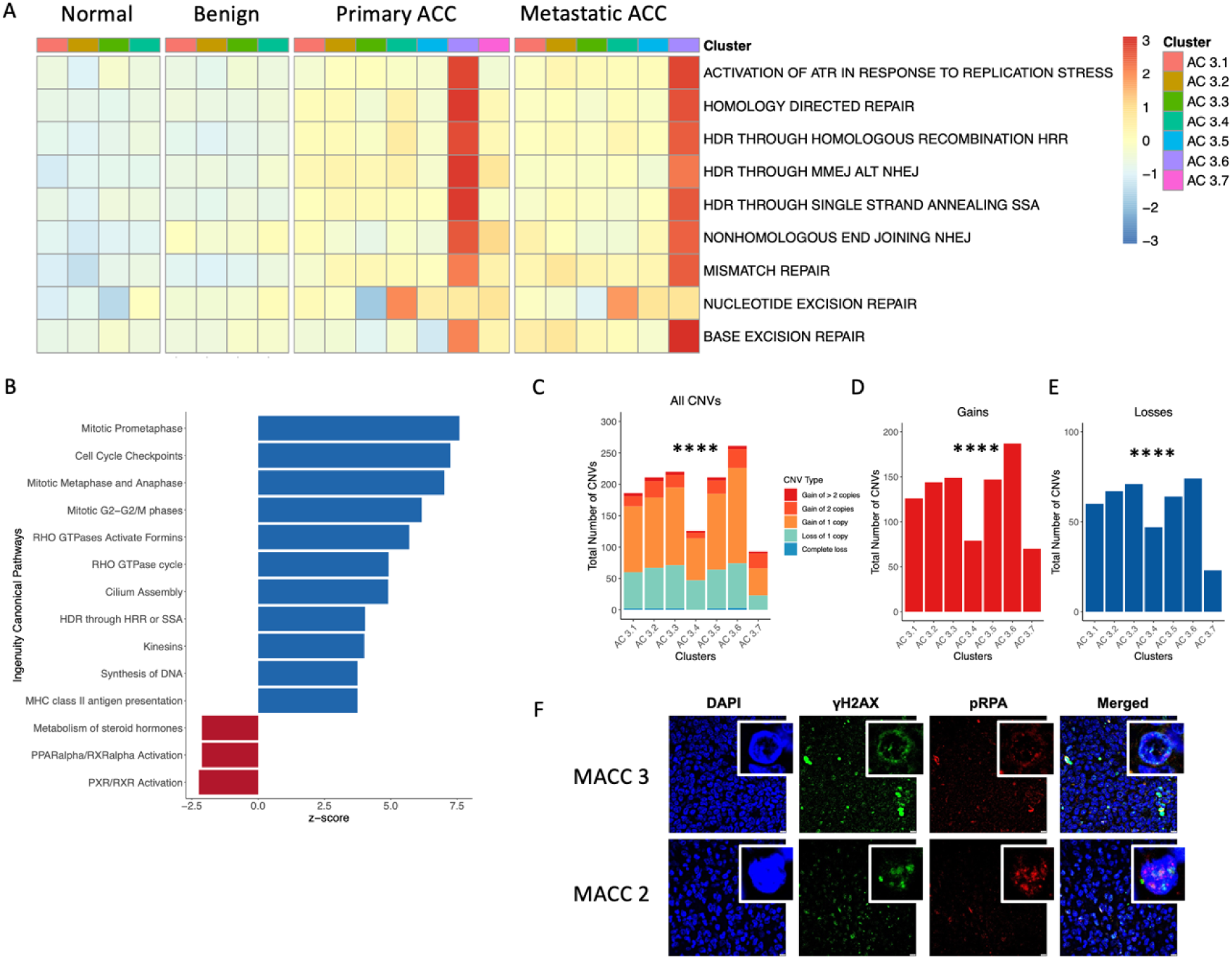
DNA Repair and Replication Pathways Are Enriched in Cluster AC 3.6. A. Heatmap demonstrating average module scores for REACTOME replication stress and DNA repair pathways downloaded from MSigDB. Clusters with fewer than 50 cells were excluded from the analysis. In the heatmap, average module scores are scaled per row. B. Bar plot demonstrating Ingenuity Pathway Analysis pathways upregulated and downregulated in cluster AC 3.6. Pathways are ordered by the z-score, and top 10 upregulated and top 3 downregulated pathways are displayed. Only 3 pathways were found to be downregulated in cluster AC 3.6. C. Bar plot showing total numbers of copy number variations (CNVs) inferred by InferCNV on a per-cluster basis. Chi-square test was used to calculate statistical significance (**** - FDR <= 0.0001). D and E. Bar plots showing the total number of gains (D) and losses (E) inferred by inferCNV on a per-cluster basis. For both plots, chi-square test was used to calculate statistical significance (**** - FDR <= 0.0001). F. Representative immunofluorescence images of formalin fixed paraffin embedded (FFPE) tissues from two patients stained for γH2AX (green), pRPA (red), and DAPI nuclear stain (blue). Scale bars, 10 µm. Zoom-in panels show details of nuclei, gH2AX foci and pRPA foci.

### ATR-dependent replication stress in ACC

To assess for the presence of replication stress and DNA damage in ACC, we performed immunofluorescence on patient specimens (MACC 2 and MACC 3) for the DNA damage marker, □H2AX and the replication stress marker phosphorylated RPA. Both proteins localize to form foci, or clusters, in regions of DNA damage or replication stress.^65^ Both ACC specimens exhibited □H2AX and RPA foci (Figure 5F, Supplementary Figure S7, Supplementary Table S3). We next sought to evaluate DNA damage in the human NCI-H295R ACC cell line to establish an *in vitro* model for further mechanistic studies and found □H2AX foci indicating a cellular response to DNA damage in the absence of cytotoxic treatments (Figure 6A, Supplementary Figure S8).

**Figure 6.**
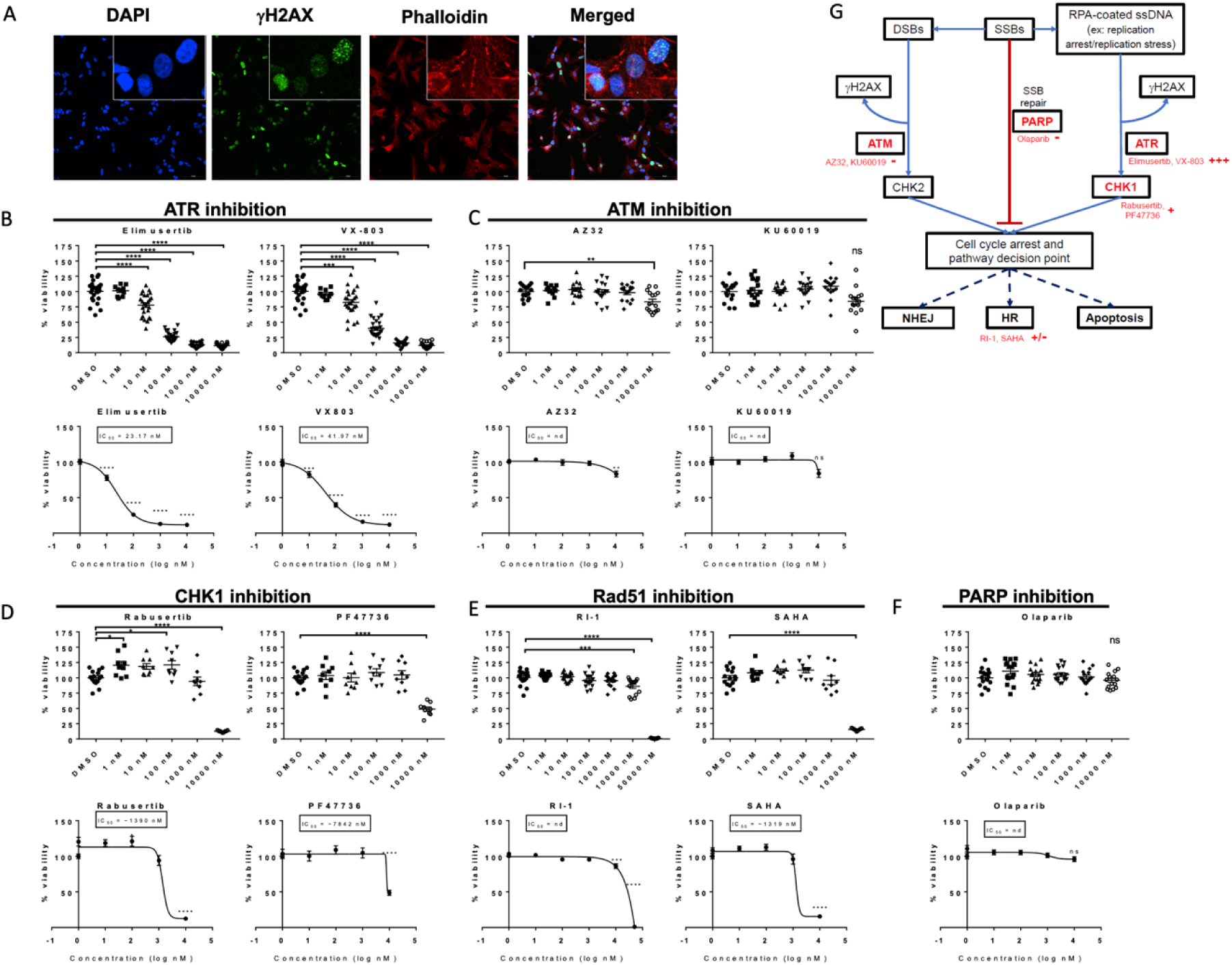
NCI-H295R Adrenocortical Cell Line Exhibits DNA Replication Stress Response. A. Immunofluorescence demonstrating that DNA damage or replication stress is present in NCI-H295R cells in the absence of cytotoxic agents as determined by the presence of intranuclear H2AX foci. In addition to the anti-H2AX antibody, the cells were stained with DAPI as nuclear stain and phalloidin. Images were obtained using a 20X objective on a Nikon^®^ AXR confocal microscope. Insets represent a cropped portion of the same fields at a 5X higher magnification. B through F. Water-soluble tetrazolium dye cell viability assays demonstrating sensitivity of the NCI-H29R5 cell line to various drugs targeting replication stress and DNA damage repair pathways: elimusertib and VX-803 (ATR inhibitors, 6B), AZ32 and KU60019 (ATM inhibitors, 6C), rabusertib and PF47736 (CHK1 inhibitors, 6D), RI-1 and SAHA (RAD51 inhibitors, 6E) and olaparib (PARP inhibitor, 6F). For each of the drug treatments, top panels display raw data and bottom panels display 4-parameter non-linear lines of best fit. G. Schematic summarizing replication stress and DNA damage repair pathways and drug treatments targeting those pathways. nd – non-determined. SAHA - suberoylanilide hydroxamic acid, ns – not significant, * - p<0.05, ** - p < 0.01, *** - p<0.001, **** - p<0.0001.

To assess for potential therapeutic vulnerabilities within the DNA damage and replication stress response in ACC, we assessed sensitivity of NCI-H295R to small molecule inhibitors of ATR kinase, elimusertib and VX-803, and of ATM kinase, AZ32 and KU60019. NCI-H295R cells were sensitive to both ATR inhibitors with IC_50_ values ranging from 23 to 42 nM (Figure 6B). In contrast, ATM inhibition with AZ32 and KU60019 had minimal cytotoxic effects, even at concentrations of 10,000 nM (Figure 6C). Prior studies have identified IC50 values ranging from 100 to 300 nM and 500 to 20,000 nM for ATR monotherapy and ATM monotherapy respectively in cell lines for other solid tumors.^66–69^ ATR is recruited to sites of replication stress and enacts downstream signaling via phosphorylation of specific targets to arrest the cell cycle and ensure repair of damaged replication forks.^70^ Therefore, we next assessed the response of NCI-H295R cells to inhibition of enzymes in the DNA repair response downstream of ATR, including checkpoint kinase 1 (CHK1, a direct target of ATR)^70^ and Rad51, a key regulator of homologous recombination that is not a direct ATR target.^71^ Inhibition of CHK1 with rabusertib or PF47736 resulted in a modest effect on viability with IC_50_ values between 1000 to 10,000 nM (Figure 6D), which is higher than previously reported IC_50_ values in cell lines considered sensitive to CHK1 monotherapy.^69,72,73^ Inhibitors of Rad51 including RI-1 (specifically inhibits Rad51) and SAHA (HDAC inhibitor which inhibits Rad51), had minimal effect on viability except at highest concentrations (50000 nM for RI-1 and 10000 nM for SAHA, Figure 6E). Finally, inhibition of poly-ADP-ribose polymerase (PARP), a DNA repair protein critical for single strand break repair and base excision repair pathways, with olaparib had no effect on NCI-H295R viability (Figure 6F). Prior studies have identified IC_50_ values for olaparib ranging from 100 to 20000 nM in sensitive cell lines.^74,75^ Together, these data indicate the presence of replication stress and specific sensitivity to ATR inhibition in an ACC cell line.

PTOs are 3-D cultures comprised of multiple cell types that can recapitulate the heterogenetic nature of the intratumoral environment and can model many aspects of tumor complexity *in vitro*.^76,77^ To further assess the role of replication stress in ACC tumorigenesis, we generated hormonally active ACC PTOs from patient 5 (**Table 1, OSU1**) and a second ACC patient with biochemical evidence of hypercortisolism (**OSU2**). Neither patient received neoadjuvant cytotoxic therapies prior to surgery. Both ACC PTO lines proliferated long-term in culture, expressed high levels of the adrenal cortex marker SF1, and produced cortisol (Figures 7A-C). Similar to the NCI-H295R cell line, both untreated ACC PTO lines demonstrated LH2AX foci (Figure 7B, Supplementary figure S9). To determine if ACC PTOs exhibit sensitivity to ATR inhibition, we treated the ACC PTOs with elimusertib and VX-803, which reduced viability at IC_50_ values of 139 to 326 nM and 2000 to 3000 nM, respectively (Figure 7D). We also treated the ACC PTOs with the ATM inhibitors, AZ32 and KU60019, neither of which caused any appreciable reduction in viability (Figure 7D). These findings reinforce a critical role of ATR in ACC and further indicate a potential role for ATR inhibition in treatment of ACC.

**Figure 7.**
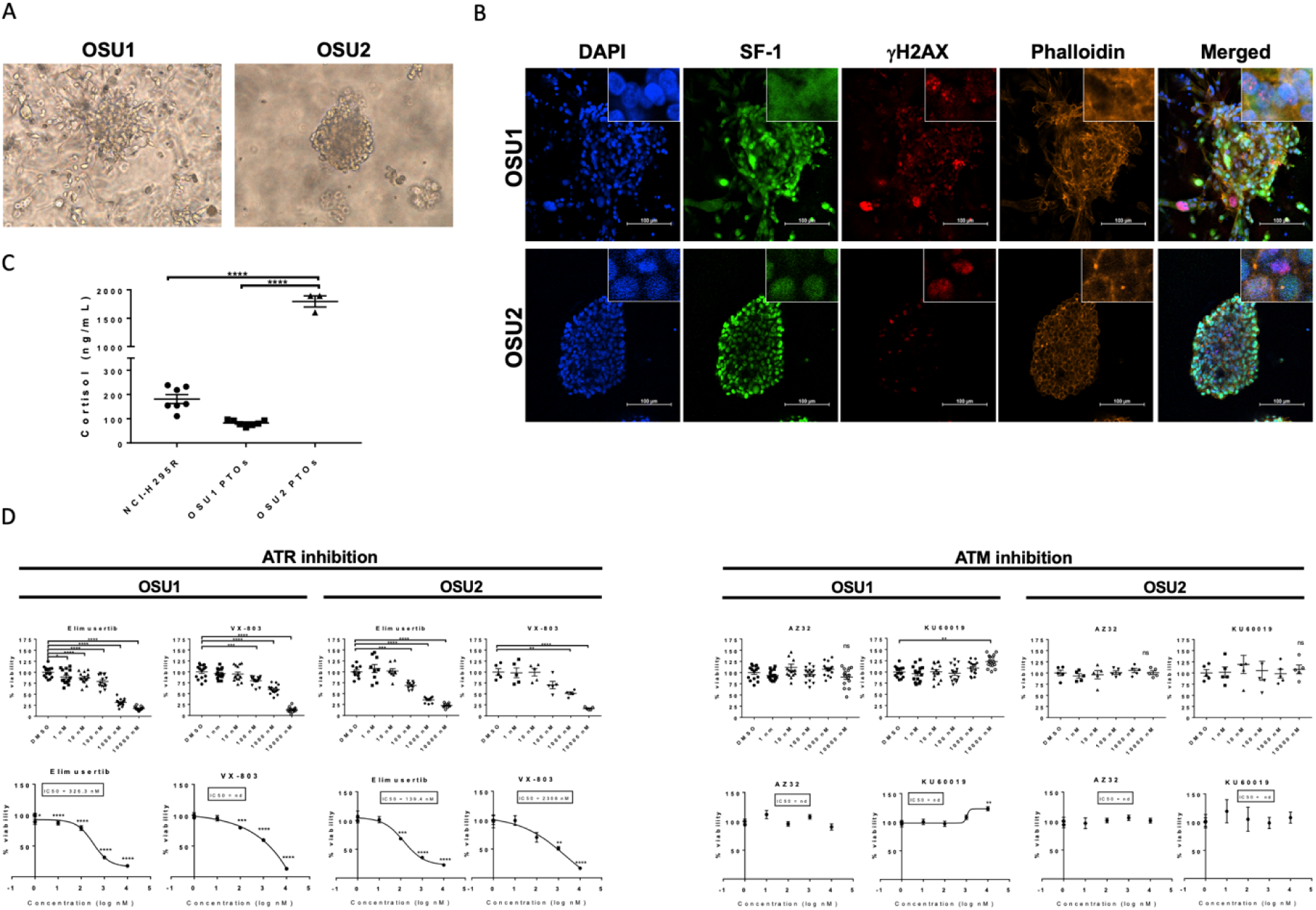
ACC PTOs exhibit DNA Replication Stress Response. A. ACC PTOs proliferate in culture long-term. Brightfield images of two ACC PTO lines (**OSU1** and **OSU2**) were taken using a 20X objective >6 months after original tumor cell isolation. B. Immunofluorescence demonstrating that ACC PTOs express steroidogenic factor 1 (SF-1) and also showing that DNA damage or replication stress is present in ACC PTOs in the absence of cytotoxic agents as determined by the presence of intranuclear LH2AX foci. In addition to the anti-H2AX antibody and anti-SF-1 antibody, the cells were stained with DAPI as nuclear stain and phalloidin. Images were taken using a 20X objective on a Nikon^®^ AXR confocal microscope. Insets represent a cropped portion of the same fields at a 5X higher magnification. C. ELISA demonstrates that ACC PTOs secrete cortisol. D. Water-soluble tetrazolium dye cell viability assays demonstrate that ACC PTOs exhibit sensitivity to ATR but not ATM inhibition. For each of the drug treatments, left panel displays raw data and right panel displays 4-parameter non-linear lines of best fit. nd - not determined, ns – not significant, * - p<0.05, ** - p < 0.01, *** - p<0.001, **** - p<0.0001.

## DISCUSSION

In this study, we use snRNA sequencing to demonstrate changes in immune, fibroblast and adrenal cortex landscape in PACC and MACC specimens relative to non-malignant adrenal cortex tissues. As such, we have identified several potential therapeutic approaches for further investigation in ACC clinical management, as outlined below. Immunotherapy options for ACC have been characterized by poor responses.^7^ Our data indicate that macrophages are the dominant immune cell and that SPP1+ macrophages are increased in ACC. Recent studies demonstrate that SPP1+ macrophages are associated with tumor progression and worse clinical outcomes.^32,33,36,78,79^ In addition, ACC macrophages expressed markers of immunosuppression. Collectively, these data suggest targeting macrophages with antibodies or small molecules, such as blockade of the CD47-SIRPα axis or anti-PDL1 antibodies^38^, to induce activation and tumor cell phagocytosis may be a viable immunotherapy strategy in ACC. In contrast, we found lymphoid infiltration only in the absence of excess cortisol production, suggesting that immunotherapy alone may be more effective in non-functional ACCs and that concurrent cortisol blockade may improve effectiveness of immunotherapy in the presence of hypercortisolism.

Interestingly, our analysis of fibroblasts showed a loss of fibroblast subpopulations in ACC. In addition, the same fibroblast population was present in primary and metastatic ACC. These findings suggest that this fibroblast subpopulation may migrate from the primary tumor to the site of metastasis and serve as a premetastatic niche to support tumor cells as in other cancers.^80^ Fibroblasts in ACC expressed increased *FAP* and *PDGFRB*. Targeting FAP and PDGFRB resulted in tumor reduction in preclinical studies of other cancers.^81–83^ Further studies are necessary to elucidate the role of fibroblasts in ACC tumor progression and identify potential therapeutic targets.

We found that ACC was characterized by a unique adrenal cortex subpopulation defined by a signature for replication stress and DNA damage response pathways. This subpopulation may correspond to the ACC M subpopulation identified by Tourigny et al.^16^ Although Tourigny et al. identified a signature for mitosis in the ACC M subpopulation, they did not identify a signature for replication stress or DNA repair. Our extended findings may be attributed to differences in sequencing methodologies and patient population. While Tourigny et al.^16^ predominantly included early-stage tumors, which can provide insight into the developmental origin of ACC, our study included stage 4 tumors, which may provide insights for developing targeted therapies. Other studies have established that genes implicated in DNA damage and cell cycle regulation are overexpressed in ACC and are associated with worse overall survival in patients.^8,84^ In addition, Gara et al. have shown that *APOBEC3B,* which is known to induce DNA damage, is overexpressed in ACC tumors with *TP53* mutations.^85^ Prior work has also shown that metastatic ACC exhibits a higher mutation burden compared to primary tumors,^86^ indicative of genomic instability. In addition to identifying replication stress signature using snRNAseq data, our work mechanistically evaluated these pathways in ACC.

Cancer cells experience chronic replication stress due to oncogene activation and sustained proliferative signaling. Furthermore, many chemotherapeutic agents induce replication stress by depleting nucleoside pools required for DNA replication or causing direct DNA damage in tumor cells.^88^ ATR is a primary upstream orchestrator of the replications stress response pathway. By inducing G2-S phase cell arrest and DNA repair as well as replication fork stabilization, ATR allows cells to adapt to and prevent replication-stress induced cell death.^70,89,90^

Our findings suggest that ATR can be therapeutically targeted in ACC irrespective of cortisol secretion. ATR inhibitors drive cells with increased DNA damage to prematurely enter mitosis, resulting in cell death. In addition to preclinical studies, some ATR inhibitors have also been investigated in clinical trials as single agents or in conjunction with other therapeutic modalities, such as radiotherapy or chemotherapy. In advanced solid tumors, ATR inhibitors as monotherapy are well-tolerated and have resulted in partial or complete response.^93,94^ Other preclinical studies and clinical trials show ATR inhibitors and chemotherapeutic agents can work synergistically to show efficacy in triple negative breast cancer, melanoma, platinum-resistant high-grade serous ovarian cancer, and other advanced solid tumors.^91,94–99^ Thus, by demonstrating ATR-dependent replication stress in ACC, we identify a promising novel treatment strategy for this deadly disease.

Identifying biomarkers to predict ATR sensitivity in ACC can improve potential efficacy. In non-small cell lung cancer (NSCLC), loss of BRG1, a chromatin remodeling enzyme, was identified as a biomarker for efficacy of ATR inhibition.^100^ Loss of ARID1A, genomic instability, ATM/G1 pathway abnormalities, and high tumor inflammation have also been identified as potential biomarkers of sensitivity to ATR inhibition.^92^ Although some preclinical models, including chronic lymphocytic leukemia, have demonstrated ATR sensitivity in conjunction with *TP53* mutation, this relationship was not seen in the aforementioned clinical trial of ATR inhibition in solid tumors.^102–104^ Importantly, *TP53* inactivating mutations are present in 21-33% of ACC,^8,105^ so future work elucidating the role of p53 inactivation or other biomarkers in the context of ATR sensitivity are necessary.

CHK1 is a target of ATR that plays a prominent role in the ATR signaling pathway by activating checkpoints, delaying cell cycle progression, and enabling DNA repair to proceed.^89,106^ Our data show that the NCI-H295R cell line and ACC PTOs exhibit notable sensitivity to ATR inhibition and modest sensitivity to CHK1 inhibition. These data suggest that ATR-dependence in ACC may in part occur via alternative downstream targets, such as SMARCAL1, which functions to promote replication fork stability and fork restart.^107^ Additional non-CHK1 targets of ATR, which are implicated in the replication stress response, include WRN and FANCI.^108,109^ Further studies are needed to determine if these targets play a role in ACC tumor progression.

Taken together, our results demonstrate diversity of transcriptional signatures in ACC and implicate several possible strategies for targeted treatment of ACC. Moreover, our results exemplify how elucidation of cellular heterogeneity combined with mechanistic evaluation in PTOs might direct therapeutic decisions and improve patient outcomes in ACC and other rare cancers.

## METHODS

### Sample collection

Patients were consented for collection of specimens and clinical information using a study protocol approved by The Ohio State University Institutional Review Board. Normal adrenal and adrenal adenoma (AA) specimens were collected from patients diagnosed with adrenal hypercortisolism and included one normal adrenal-adrenal adenoma pair. Primary (PACC) and metastatic ACC (MACC) samples were collected from patients diagnosed with ACC and included one tumor-metastasis pair. After collection, samples were frozen for further processing. We performed snRNAseq as our samples included previously banked frozen samples.^110^ Somatic and germline mutations are reported for patients who underwent genetic testing using Tempus (Chicago, IL), Foundation Medicine (Boston, MA), or CustomNext-Cancer testing from Ambry Genetics (Aliso Viejo, CA). Sample details are summarized in Table 1.

### Single nuclei RNA sequencing

#### Nuclei isolation and library preparation

Nuclei were isolated from frozen tissue as described in Lacar et al. and sorted using the Bigfoot Spectral Cell Sorter (Thermo Fisher Scientific, Waltham, MA). For each sample, 16,000 nuclei were sorted directly into the 10X Genomics master mix to load approximately 10,000 nuclei into a 10x Genomics Chromium device for microfluidic-based capture of single nuclei.^111^ Reverse transcription, cDNA amplification and library preparation were performed according to the manufacturer’s protocol for Chromium Next GEM Single Cell 3’ Gene Expression kit. Resulting libraries were sequenced on an Illumina NovaSeq 6000 instrument to generate paired-end sequencing data.

#### Data preprocessing

Fastq files were generated using the mkfastq command from 10x Genomic CellRanger. Alignment to the GRCh38 human reference, filtering, barcode counting, and UMI counting steps were performed using the 10x Genomics CellRanger software suite v7.0 following the default parameters.

#### Quality control, filtering, and doublet removal

Downstream analyses were performed using Seurat v4 for R 4,1,1 and 4.2.2.^19^ For quality control purposes, individual objects were filtered to remove nuclei that contained more than 5% of mitochondrial transcripts. In addition, data were filtered to only include nuclei that contained a minimum of 200 detected genes for all the samples and a maximum of 8000 genes (for Normal and AA 1 samples), a maximum of 7000 genes (for AA 2 sample), a maximum of 13000 genes (for PACC 1 sample), a maximum of 12000 genes (for PACC 2 sample), a maximum of 9000 genes (for PACC 3, PACC 4, and MACC 4 samples), a maximum of 15000 genes (for MACC 1 sample), a maximum of 11500 genes (for MACC 2 sample), and a maximum of 14000 (for MACC 3 sample). Upper filtering cutoffs were selected based on differences in data distribution between samples. As a result of filtering, the following numbers of nuclei were excluded from further analysis: 17 nuclei (Normal sample), 49 nuclei (AA 1 and MACC 1 samples), 29 nuclei (AA 2 sample), 70 nuclei (PACC 1 sample), 381 nuclei (PACC 2 sample), 328 nuclei (PACC 3) sample, 171 nuclei (PACC 4 sample), 31 nuclei (MACC 2 sample), 553 nuclei (MACC 3 sample), and 5 nuclei (MACC 4 sample). Doublets were detected and removed from individual objects using the DoubletFinder package v2.0 for R.^112^

#### Normalization and cell cycle regression

To control for cell-cycle effects on transcriptional programs, we performed cell cycle regression. After DoubletFinder, cell cycle scores were calculated for the individual objects using the CellCycleScoring() function based on the list of cell cycle marker genes published in Tirosh et al.^113^ Normalization and variance stabilization were performed on the individual objects using the SCTransform() function from the sctransform package v0.3 for R with “vst.flavor” argument set to v2 and “vars.to.regress” argument set to regress the percentage of mitochondrial content, G2/M and S cell cycle scores.^114^

#### Data integration

To minimize patient-to-patient variability, samples were integrated using the RPCA algorithm in Seurat.^115^

#### Dimensionality reduction and clustering

Principal component analysis (PCA) was performed on the integrated Seurat object. The first 30 principal components were used for dimensionality reduction via the Uniform Manifold Approximation and Projection (UMAP) method. Clusters were defined using the FindClusters() function for a series of resolutions between 0.1 and 1. Optimal clustering resolution of 0.1 was selected based on cluster stability, as in Long et al., using the clustree package v0.5 in _R.116,117_

#### Cell type annotation

FindAllMarkers() function in Seurat was used to define marker genes for further cluster annotation. Dot plots were generated using the DotPlot() function in Seurat with the argument “scale” set to the default value. To complement the FindAllMarkers() approach and annotate adrenal cortex clusters more precisely, we used the AddModuleScore() function in Seurat to calculate module scores based on the expression of *NR5A1* (which encodes the protein steroidogenic factor 1, SF1), *INHA*, and *MLANA*, which are all adrenal cortex biomarkers.^17,18^ Adrenal cortex clusters were characterized by higher adrenal cortex biomarker module scores when compared to lymphocyte, myeloid cell, endothelial, and fibroblast clusters.

#### Subclustering of immune cells

Myeloid- and lymphocyte-typed clusters were subset from the initial integrated Seurat object and re-processed using SCTransform as described above. Samples were then merged and re-integrated using Harmony^26^ at a resolution of 0.2. Dimensionality reduction and clustering steps were followed as described above. This process was repeated after non-immune cells were subset out. ggplot2^118^ was used to generate a histogram of immunocyte proportions and the FindAllMarkers() function in Seurat was used to define marker genes for cluster annotations displayed using the DotPlot() function. The macrophage-typed clusters were then further subset, re-processed, and re-integrated using Harmony at a resolution of 0.2. This process was repeated after excluding a small number of CD3-expressing lymphocytes. Dimensionality reduction, clustering steps, and cell type annotation were performed as described above. ggplot2^118^ was used to generate a histogram of macrophage subtypes for each sample type (normal adrenal, AA, PACC, MACC). The function propeller() from the speckle^119^ package for R was used to compare subtype proportions across sample types. False discovery rate (FDR) is reported. Average gene expression was calculated across each sample type and scaled average expression was displayed in a heatmap using the DoHeatmap() function. There were not enough cells for the CD8+ T cell, CD4+ T cell, and natural killer (NK) cell clusters to subset and reprocess, re-integrate, and recluster. Therefore, these lymphocytes were subset from the macrophage and leukocyte re-integrated object and average gene expression was calculated across each sample type and displayed in a heatmap as described above.

#### Subclustering of fibroblasts

The fibroblast cluster was subset from the initial integrated Seurat object, reprocessed, and re-integrated using Harmony to increase clustering resolution. After dimensionality reduction and clustering, optimal resolution of 0.2 was selected as described above. As a result, 10 clusters were identified. Four clusters including adipocytes were excluded, after which the fibroblasts were reprocessed and re-integrating using Harmony. PACC 2 was excluded from analysis due to having only 5 fibroblasts. After dimensionality reduction and clustering, optimal resolution of 0.1 was selected as described above. As a result, 5 clusters of fibroblasts were identified. UMAP projection was displayed using the DimPlot() function in Seurat. Marker genes were identified using the FindAllMarkers() function and visualized using the DotPlot() function in Seurat. Average gene expression was calculated across sample types and displayed in a heatmap as described above.

#### Subclustering of adrenal cortex cells

To increase clustering resolution, adrenal cortex cells were subset from the integrated Seurat object and reprocessed using normalization, cell-cycle regression, integration using RPCA, dimensionality reduction, and clustering steps as in processing the initial Seurat object. Optimal clustering resolution of 0.1 was selected as described above. FindAllMarkers() function was used to identify marker genes for each of the clusters. Marker genes were visualized using the DotPlot() function as described above. In addition, adrenal cortex module scores were calculated for each of the clusters as described above. As a result, 6 adrenal cortex clusters and 1 cluster of adrenal medulla were identified. To further increase clustering resolution, the adrenal medulla cluster was excluded, and adrenal cortex cells were again subset from the integrated Seurat object and reprocessed using normalization, cell-cycle regression, integration, dimensionality reduction, and clustering steps as described above. Optimal resolution of 0.1 was selected as described above. As a result, 7 clusters of adrenal cortex cells were identified; these cluster assignments were used for all downstream analyses.

#### Cell type proportions

To compare cell type proportions of adrenal cortex cells across the samples, we used the propeller method from the speckle package v0.0.3 in R.^119^

#### Pathway analysis

To characterize adrenal cortex clusters based on gene expression patterns, we used the AddModuleScore() function from the Seurat package to calculate module scores using a selection of Hallmark pathways downloaded from MSigDB using the msigdbr package v7.5 in R.^59,60^ We further calculated average module scores for each of the clusters to generate the heatmap for Figure 4C. We also calculated average module scores for each of the clusters on a per-tumor type basis to generate the heatmap for Supplementary Figure S4. We performed a similar analysis using a selection of replication stress and DNA damage response REACTOME pathways downloaded from MsigDB^59–61^ We calculated module scores as in Wu et al.^35^ for each of the selected REACTOME pathways, and then calculated average module scores for each cluster on a per-tumor type basis (for Figure 5A) and on a per-sample basis (for supplementary Figure S5), which were displayed on a heatmap. To generate heatmaps, we used the pheatmap() function from the pheatmap package v.1.0 in R with the argument “scale” set to “row”.^120^ For all heatmaps, clusters with fewer than 50 cells were excluded from analysis. Additionally, we used QIAGEN Ingenuity Pathways Analysis (IPA) (QIAGEN Inc., https://digitalinsights.qiagen.com/IPD) to carry out pathway enrichment analysis for Figure 5B.^62^

#### InferCNV

We used inferCNV^64^ v1.8 to infer copy number alterations in the tumor samples. We used the HMM_CNV_predictions.HMMi6.hmm_mode-samples.Pnorm_0.5.pred_cnv_regions.datfile to quantify the number of chromosomal aberrations per sample. To compare the number of chromosomal aberrations between clusters, we used the chisq.test() function in R.

#### Data and code availability

snRNAseq data generated for this study has been deposited to dbGAP, accession phs003764. Code that was used to analyze the snRNAseq data is available upon request.

### Immunofluorescence of ACC

Hematoxylin and eosin–stained slides of formalin fixed paraffin embedded (FFPE) samples were screened for diagnosis, cellularity, and necrosis by a board-certified pathologist. FFPE blocks were cut into 4-μm sections and deparaffinized in organic solvents. Slides were dehydrated, submerged in DAKO Antigen Retrieval Buffer (pH 9.0) (Agilent Technologies, Santa Clara, CA), and incubated at 110°C in a steam rice cooker for 30 minutes. The slides were then cooled on ice for 30 minutes, followed by 5 minutes in distilled water. As previously described^121^, samples were permeabilized with DAKO Wash Buffer (Agilent Technologies), then blocked with blocking buffer (DAKO Wash buffer; 1% BSA). Sections were stained with mouse anti-LH2AX (Millipore-Sigma, St.Louis, MO, 05-636) at 1:500 and rabbit anti-RPA (Bethyl Laboratories, Montgomery, TX, A300-245A-M) at 1:200. Secondary incubation was performed with goat anti-mouse Alexa fluor 488 and goat anti-rabbit Alexa fluor 647. DAPI (Millipore-Sigma) was added, and the slides were dehydrated with increasing concentrations of ethanol. Samples were then mounted with ProLong Gold anti-fade reagent (Thermo Fisher Scientific) and stored at –20°C. Stained samples were imaged on a Leica Thunder Imager Microscope.

### Cell culture

#### NCI-H295R

The NCI-H295R ACC cell line was originally purchased from American Type Culture Collection (ATCC, Manassas, VA). NCI-H295R cells were grown in DMEM/F12 with L-glutamine and 15 mM HEPES without phenol red (Thermo Fisher Scientific) supplemented with 2.5% Nu-Serum™ (Corning, Corning, NY), 1% penicillin-streptomycin (Sigma-Millipore), and 1x ITS Premix Universal Cell Culture Supplement (Corning). Cells were split approximately once per week and all experiments utilized cells below 20 passages.

#### ACC patient-derived tumor organoid (PTO) generation

PTOs were generated from a tumor harvested from a patient diagnosed with ACC (patient 5 in Table 1). Briefly, tumor tissue was minced then digested with 0.5 mg/mL collagenase II (Worthington Biochemical, Freehold, NJ) in DMEM/F12 (Cytiva, Marlborough, MA). Tumor cells were then seeded at 5-10×10^4^ cells per well in 24-well plates in 80% Matrigel^®^ (Corning), with DMEM/F12 supplemented with 10 nM HEPES (Cytiva), 1x Glutamax, 1x B27 Supplement (both Thermo Fisher Scientific), 1.25 mM n-acetylcysteine, 10 mM nicotinamide, 10 nM [Leu15]-Gastrin, 0.5 μM A-83-01, 1 μM PGE2 (all Sigma-Millipore), 0.5 μg/mL R-Spondin-1, 50 ng/mL EGF (both R&D Systems, Minneapolis, MN), 0.1 μg/mL FGF-10, 30 ng/mL recombinant murine Wnt3a, and 0.1 μg/mL recombinant human Noggin (all Thermo Fisher Scientific). PTOs were routinely passaged upon confluency. Brightfield images of organoid colonies were captured using an EVOS™ M5000 microscope (Thermo Fisher Scientific).

### Immunofluorescence of cultured cells and ACC PTOs

NCI-H295R cells were seeded at 1.5 x 10^5^ cells per cm^2^ on top of 12 mm diameter poly-L-lysine coated glass coverslips (Electron Microscopy Sciences, Hatfield, PA) placed in wells of a 24-well cell culture plate. After 24 hours, cells were washed with PBS, fixed with 4% paraformaldehyde (Thermo Fisher Scientific), and permeabilized with 0.5% Triton X-100. After blocking with 10% normal goat serum (Abcam, Cambridge, UK) in 0.1% Triton X-100, cells were stained with mouse anti-γH2AX (Cell Signaling Technologies, Danvers, MA, CST#80312) at 1:250 followed by secondary incubation with goat anti-mouse Alexa Fluor^®^ 488 antibody at 1:500 (Abcam #ab150117) and phalloidin-Alexa Fluor^®^ 594 at 1:1000 (Abcam #ab176757). After mounting with Fluoroshield mounting medium with DAPI (Abcam), images were obtained on a Nikon AXR confocal microscope (Nikon, Tokyo, Japan). Cells incubated without primary antibodies were used as negative controls for NCI-H295R immunofluorescence experiments.

ACC PTOs were removed from Matrigel domes and stained in a similar manner, and then mounted on the 0.2-0.4 μm double cavity well microscope slides (Electron Microscopy ciences #71883-79) and imaged similarly to NCI-H295R cells. Primary antibodies were incubated at the following concentrations overnight at 4C: rabbit anti-SF-1 1:250 (Abcam #ab51074), mouse anti-γH2AX 1:250 (Cell Signaling Technologies CST#80312), or equivalent isotype controls (Abcam #ab172730, Cell Signaling Technologies CST#5415). The following secondary antibodies were used: goat anti-rabbit Alexa Fluor^®^ 488 1:1000 (Abcam #ab150081), goat anti-mouse Alexa Fluor^®^ 546 1:500 (Thermo Fisher Scientific #A-11003) phalloidin-Alexa Fluor^®^ 647 1:400 (Thermo Fisher Scientific #A2287). Comparable isotype antibodies (Cell Signaling Technologies #5145 and Abcam #ab172730) were used as negative controls for ACC PTO immunofluorescence experiments.

### Cytotoxicity assays

#### NCI-H295R Cytotoxicity Assays

NCI-H295R cells were seeded at 1×10^4^ cells per well in a 96-well plate and allowed to adhere for 24 hours. Media was then replaced with fresh media containing specified drugs or equivalent DMSO vehicle control (Corning). Cell viability was assessed by water-soluble tertozolium-8 (WST-8) based Cell Counting Kit-8 assay (Dojindo, Mashiki, Kumamoto, Japan) per the manufacturer’s instructions at 4 days. Absorbance readings were conducted using a SpectraMax^®^ iD5 plate reader and SoftMax^®^ Pro Software (Molecular Devices, San Jose, CA). All assays were performed in biologic triplicates with 3-5 replicates per condition. The following drugs were used at concentrations ranging from 1nM to 50uM: KU60019 (Selleck Chemicals, Houston, TX), RI-1 (Thermo Fisher Scientific), suberoylanilide hydroxamic acid (SAHA, aka vorinostat, Thermo Fisher Scientific), rabusertib (Selleck Chemicals), PF47736 (Selleck Chemicals), olaparib (Cell Signaling Technologies). Elimusertib (Selleck Chemicals), VX-803 (Selleck Chemicals), and AZ32 (Thermo Fisher Scientific). Mean viability of each drug treatment group was measured as a percentage of baseline DMSO vehicle treated cell absorbance. The half-maximal inhibitory concentration (IC_50_) for each drug was calculated by performing a four-parameter nonlinear best fit analysis of log concentration by percent viability on GraphPad Prism^®^ 7.0a software (GraphPad Software, La Jolla, CA). Differences in viability across each concentration of a particular drug were compared using one-factor ANOVA with post-hoc multiple comparison analysis by Dunnett’s test in GraphPad Prism^®^ 7.0a software (GraphPad Software). All data are reported as means ± SEM with statistical significance set as p <0.05.

#### ACC PTO cytotoxicity assays

Single-cell suspensions were generated by incubating well-established ACC organoid domes with 1 mg/mL dispase II (Thermo Fisher Scientific). 1.5×10^4^ cells per well were seeded in 96-well plates in 80% Matrigel^®^ as described above. Three days after seeding, media was replaced with media containing specified drugs or DMSO vehicle control. Organoid viability was assessed at 4 days by WST-8 assay as described above using the cell free Matrigel^®^ domes for background absorbance readings. Differences in viability across each concentration of a drug were compared using one-factor ANOVA with post-hoc multiple comparison analysis by Dunnett’s test in GraphPad Prism^®^ 7.0a software (GraphPad Software, La Jolla, CA). Each drug was tested in at least 3 independent experiments with 5 replicates per experiment.

### Cortisol ELISA

Single cell suspensions of NCI-H295R cells and ACC organoid-derived cells were generated as described above. 3×10^5^ cells were plated and medium was collected at 72-hours. Cortisol was measured by ELISA per manufacturer’s instructions (Abcam #ab108665). Each experiment was conducted in triplicate and read on BioTek Synergy H1 microplate reader using Gen5 v3.10 software (Agilent Technologies). Cortisol concentrations were compared between groups using two-tailed independent student’s *t*-test. All data are reported as means ± SEM with statistical significance set was *p* <0.05.

## AUTHOR CONTRIBUTION STATEMENT

Conceptualization: P.H.D.

Methodology: P.H.D. and K.E.M.

Investigation: L.V.P., E.A.R.G., D.M.C., J.B.N., A.R., Y.S., and E.L.

Specimen Acquisition: P.H.D., J.E.P., and B.S.M.

Specimen Pathology Review: S.S.

Formal analysis: L.V.P., E.A.R.G., D.M.C., Y.S., E.L.

Data curation: L.V.P., E.A.R.G., and K.E.M.

Funding acquisition: P.H.D.

Project administration: P.H.D.

Supervision: E.R.M., K.E.M. and P.H.D.

Validation: K.E.M. and P.H.D.

Visualization: L.V.P., P.H.D.

Writing—original draft: L.V.P., E.A.R.G., and D.M.C.

Writing—review and editing: L.V.P., E.A.R.G., D.M.C., B.S.M., M.M., K.E.M., and P.H.D.

## ACKNOWLEDGMENTS

This project was supported by Ohio Cancer Research, The Victory Over Cancer Foundation, DOD W81XWH-22-PRCRP-CDA-SO, NIH R01CA279997, and NIH R21CA277083 (PHD). LVP is supported by the Pelotonia Scholars Program. DMC is supported by training grant PF-23-1036284-01-CDP via the American Cancer Society. Any opinions, findings, and conclusions expressed in this material are those of the author(s) and do not necessarily reflect those of the NIH, Pelotonia Scholars Program, OSU, or the American Cancer Society. We acknowledge resources from the Campus Microscopy and Imaging Facility (CMIF), Biospecimen Services Shared Resource and the Microscopy Shared Resource. These shared resources are supported in part by the Cancer Center Support Grant P30 CA016058, National Cancer Institute, Bethesda, MD. We also acknowledge Matthew Ringel, MD, Gary Hammer, MD, PhD, Timothy Frankel, MD, PhD, Wayne Miles, PhD, Abby Green, MD, PhD, and Vineeth Sukrithan, MD for insightful discussions.

## SUPPLEMENTARY FIGURES

**Supplementary Figure S1.**
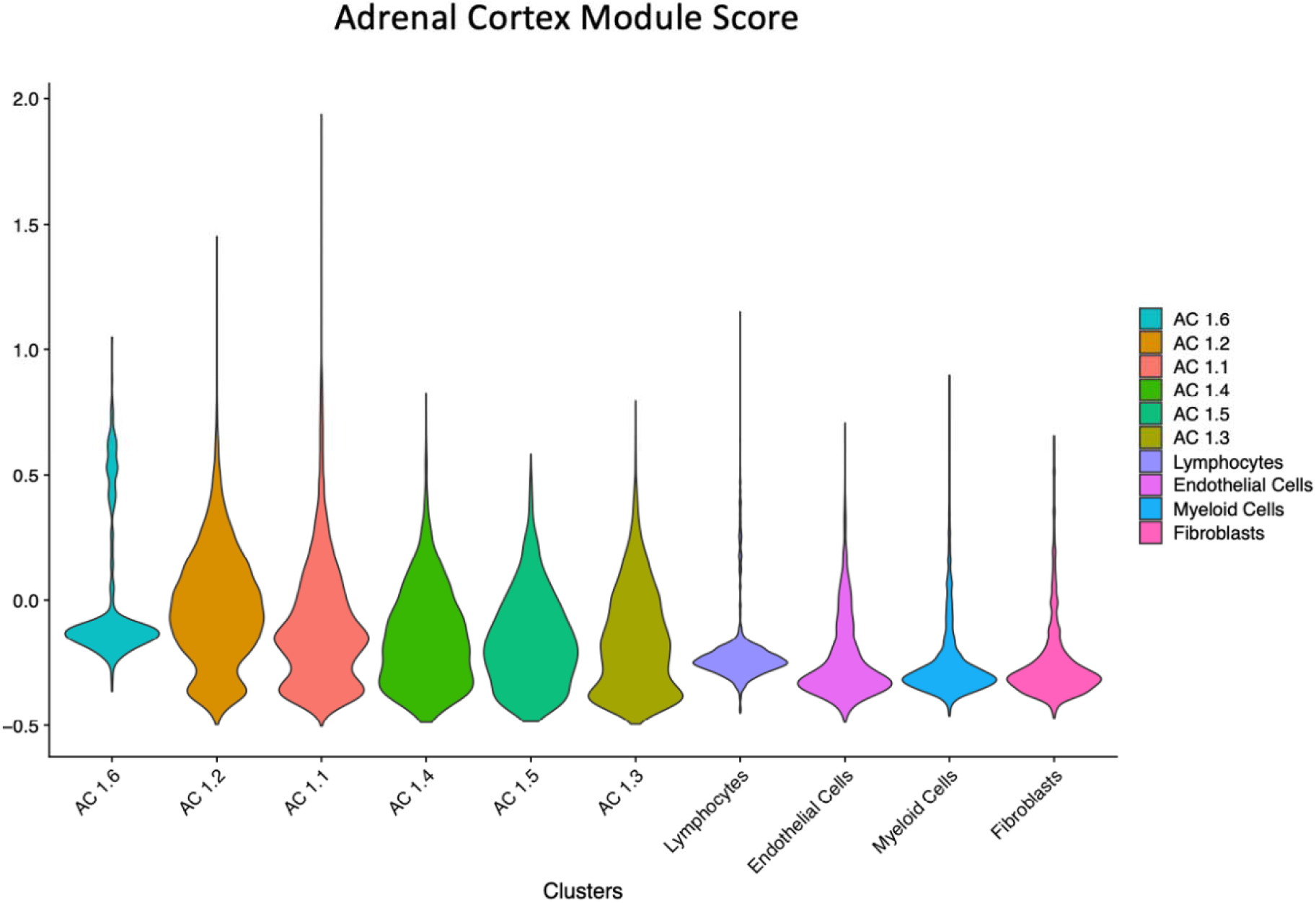
Identifying Adrenal Cortex Cells. To identify adrenal cortex clusters after initial integration, dimensionality reduction, and clustering, average module scores were calculated based on the expression of ACC markers *NR5A1*, *MLANA*, and *INHA* for each of the identified clusters. Clusters classified as adrenal cortex had higher average module scores compared to other cell types. Clusters are ordered left to right based on decreasing average module scores.

**Supplementary Figure S2.**
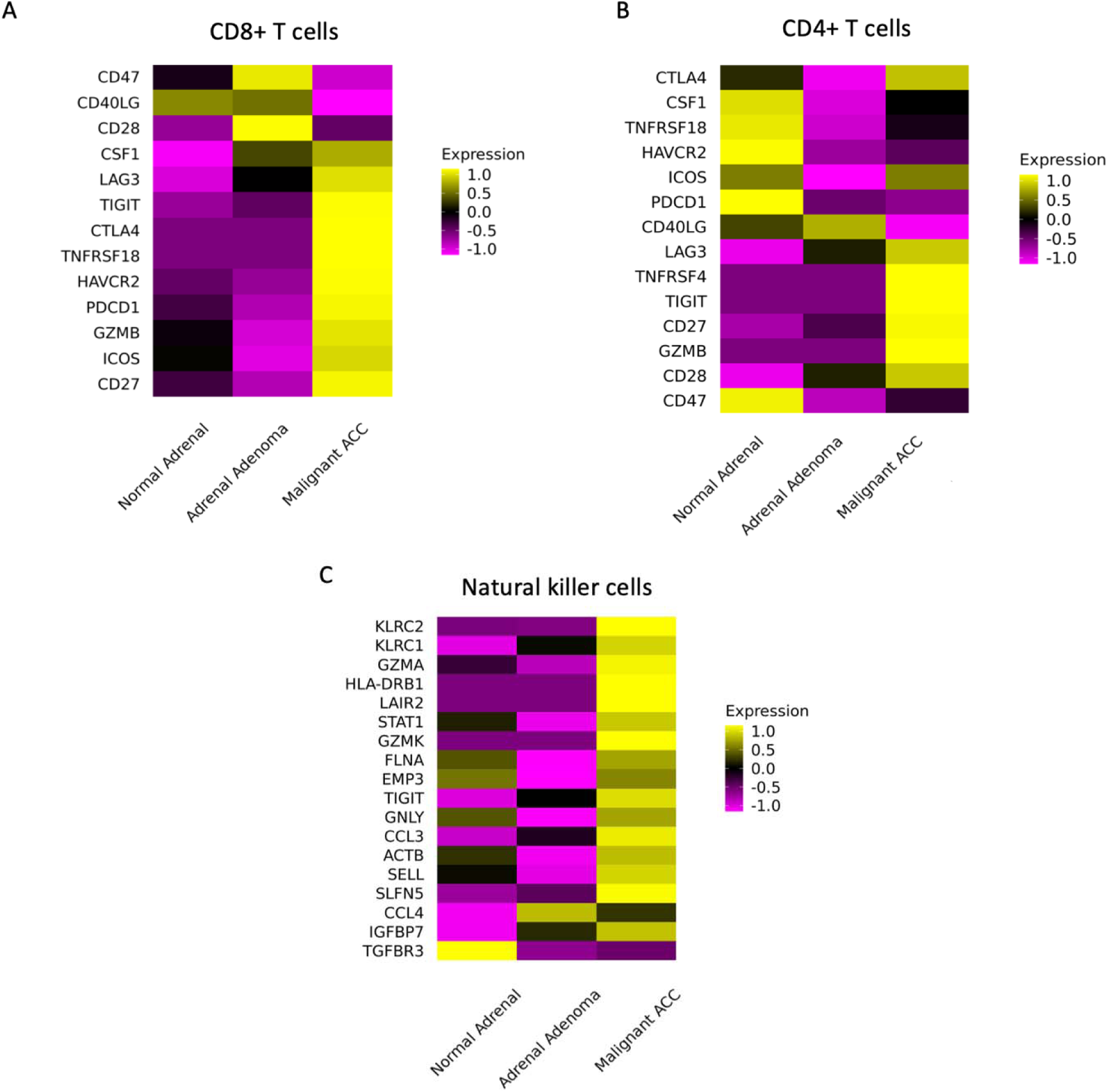
Immune Ligands Expressed by CD8+ T cells, CD4+ T cells, and Natural Killer Cells. A. Heatmaps demonstrating expression of ligands by CD8+ T cells (A), CD4+ T cells (B), and natural killer cells (C).

**Supplementary Figure S3.**
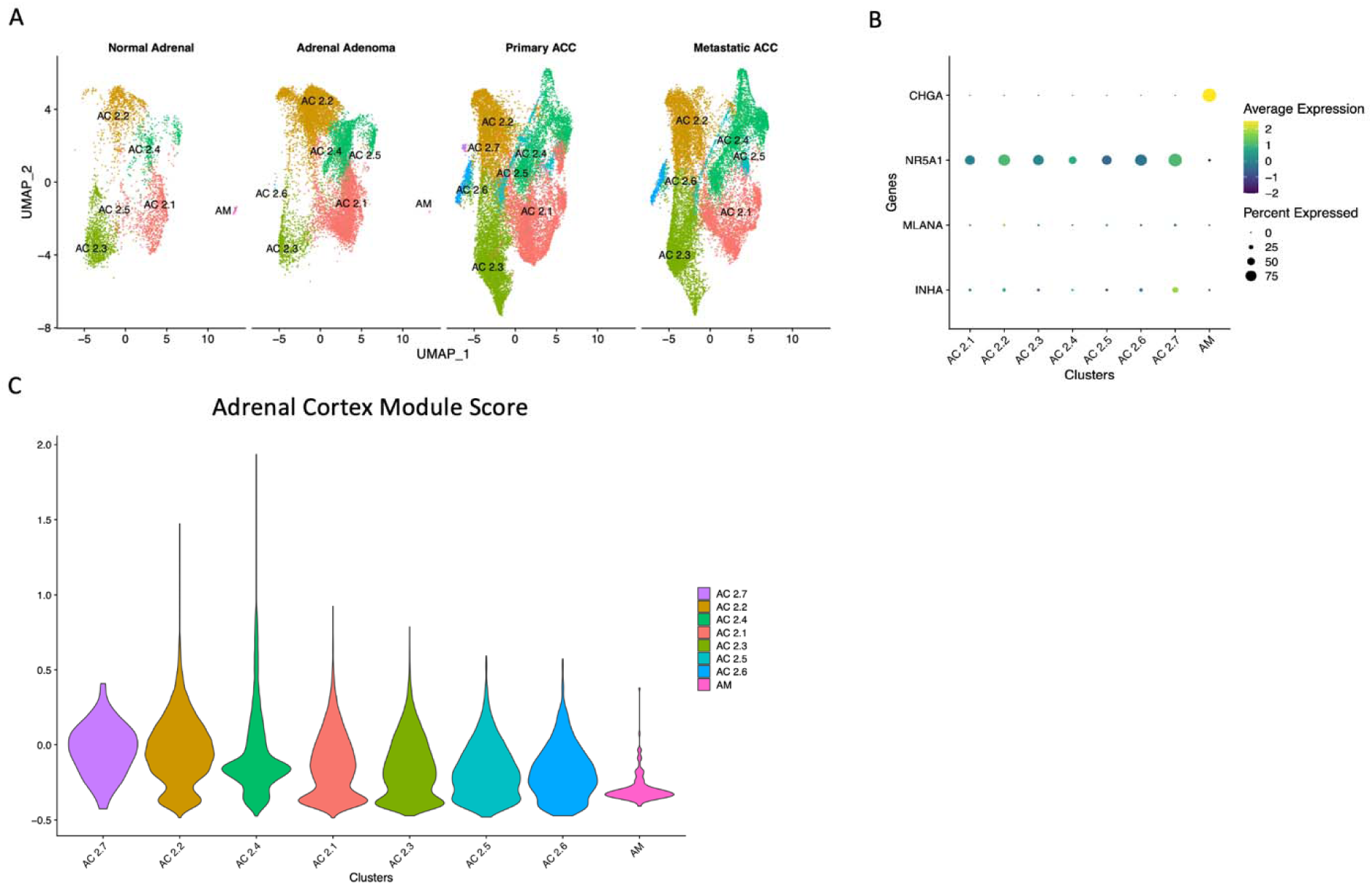
snRNAseq Demonstrates Intratumoral Heterogeneity of Adrenal Cortex Cells. A. UMAP projection showing that subclustering of adrenal cortex cells results in identification of 7 adrenal cortex clusters and 1 adrenal medulla cluster in normal adrenal, adrenal adenoma, primary ACC, and metastatic ACC specimens. B. Dotplot demonstrating marker genes that were used to annotate each of the identified clusters. C. Violin plot demonstrating average module scores calculated based on ACC marker expression as in Supplementary Figure S1. Clusters are ordered left to right based on decreasing average module scores.

**Supplementary Figure S4.**
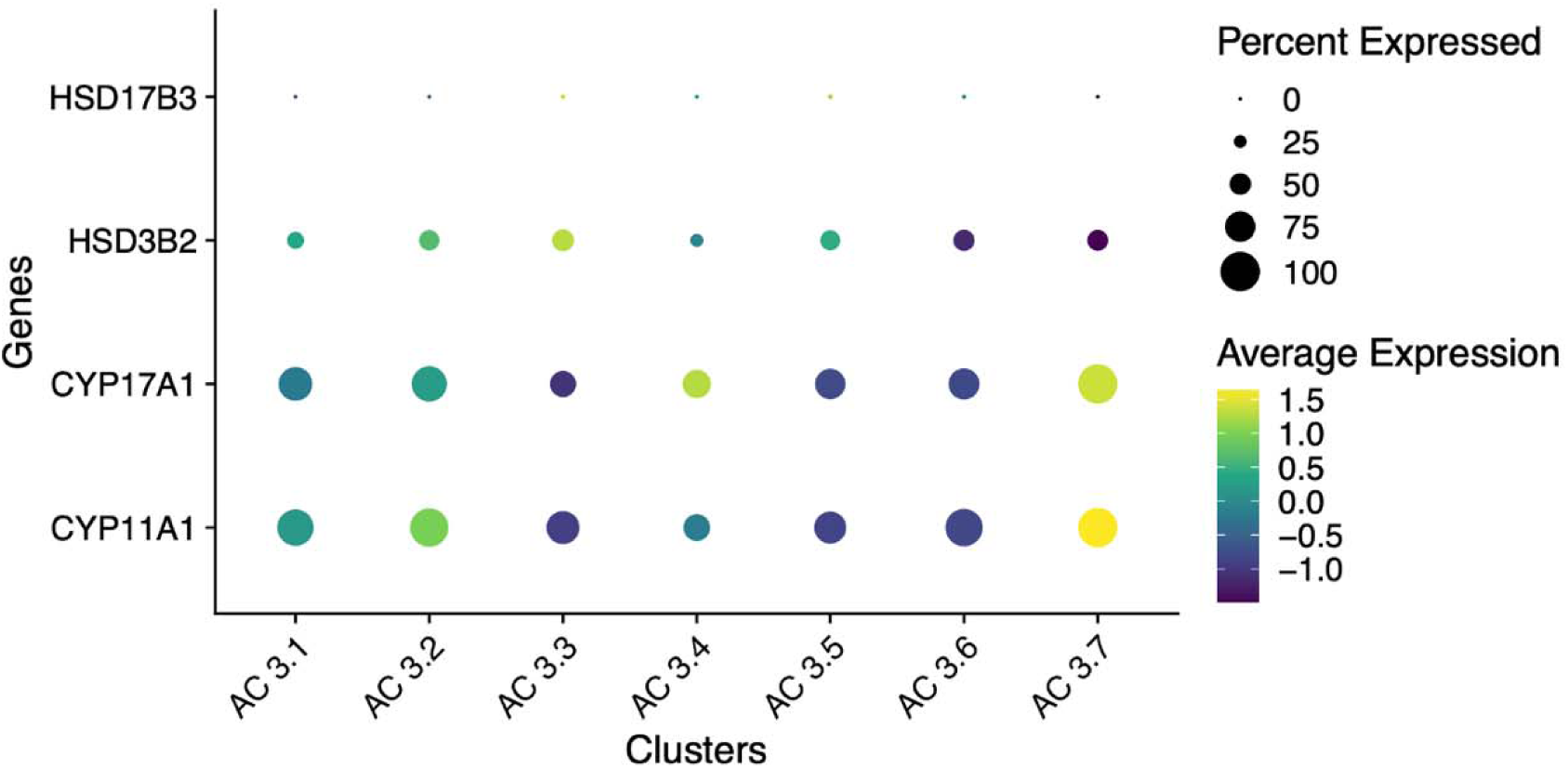
Dotplot demonstrating expression of enzymes involved in testosterone, DHEAS, and androstenedione synthesis. Cluster AC 3.7 demonstrates higher expression of *CYP11A1* and *CYP17A1*, which are involved in testosterone, DHEAS, and androstenedione production. For the dot plot, scaled average expression is displayed.

**Supplementary Figure S5.**
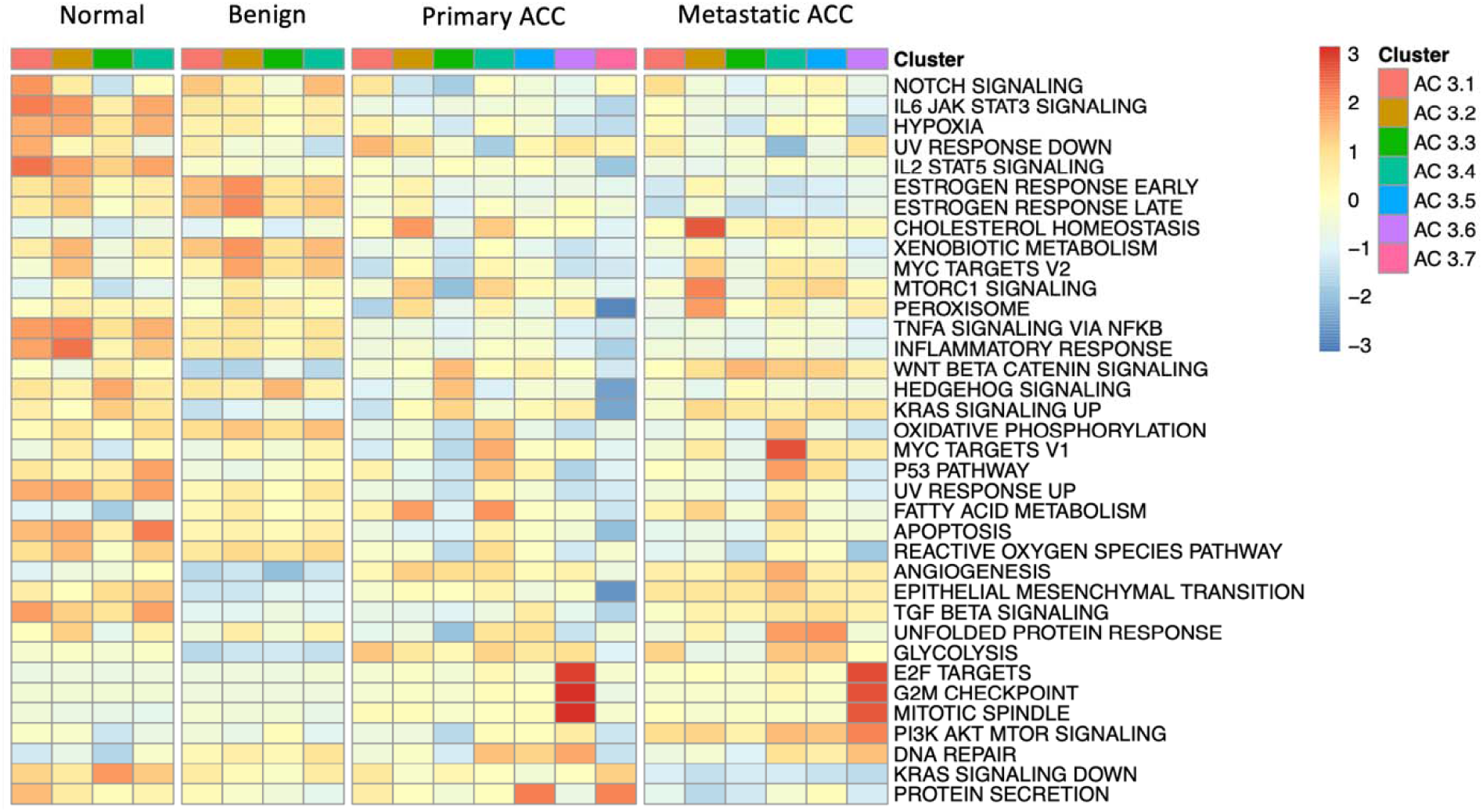
Hallmark Pathway Analysis Identifies Gene Expression Signatures on a Per-Cluster Level. Heatmap demonstrating average module scores for a selection of Hallmark pathways downloaded from MSigDB for each of the identified clusters split by sample type, expanded from Figure 2. Clusters with fewer than 50 cells were excluded from analysis. In the heatmap, average module scores are scaled on a per-row basis.

**Supplementary Figure S6.**
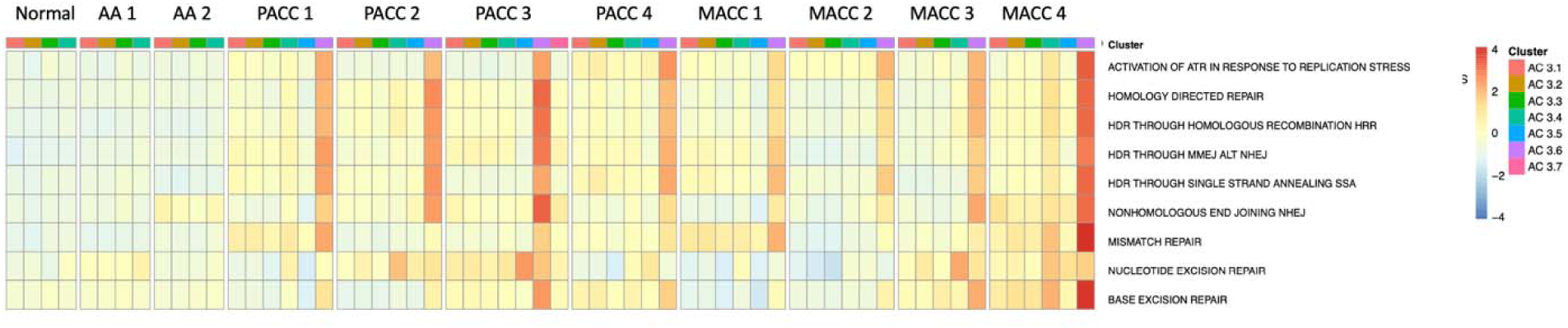
DNA Repair Pathways are Enriched in Cluster AC 3.6. A. Heatmap demonstrating average module scores for a selection of REACTOME pathways downloaded from MSigDB for all adrenal cortex clusters split by individual sample, extended from Figure 5. Clusters with fewer than 50 cells were removed from analysis. In the heatmap, average module scores are scaled on a per-row basis.

**Supplementary Figure S7.**
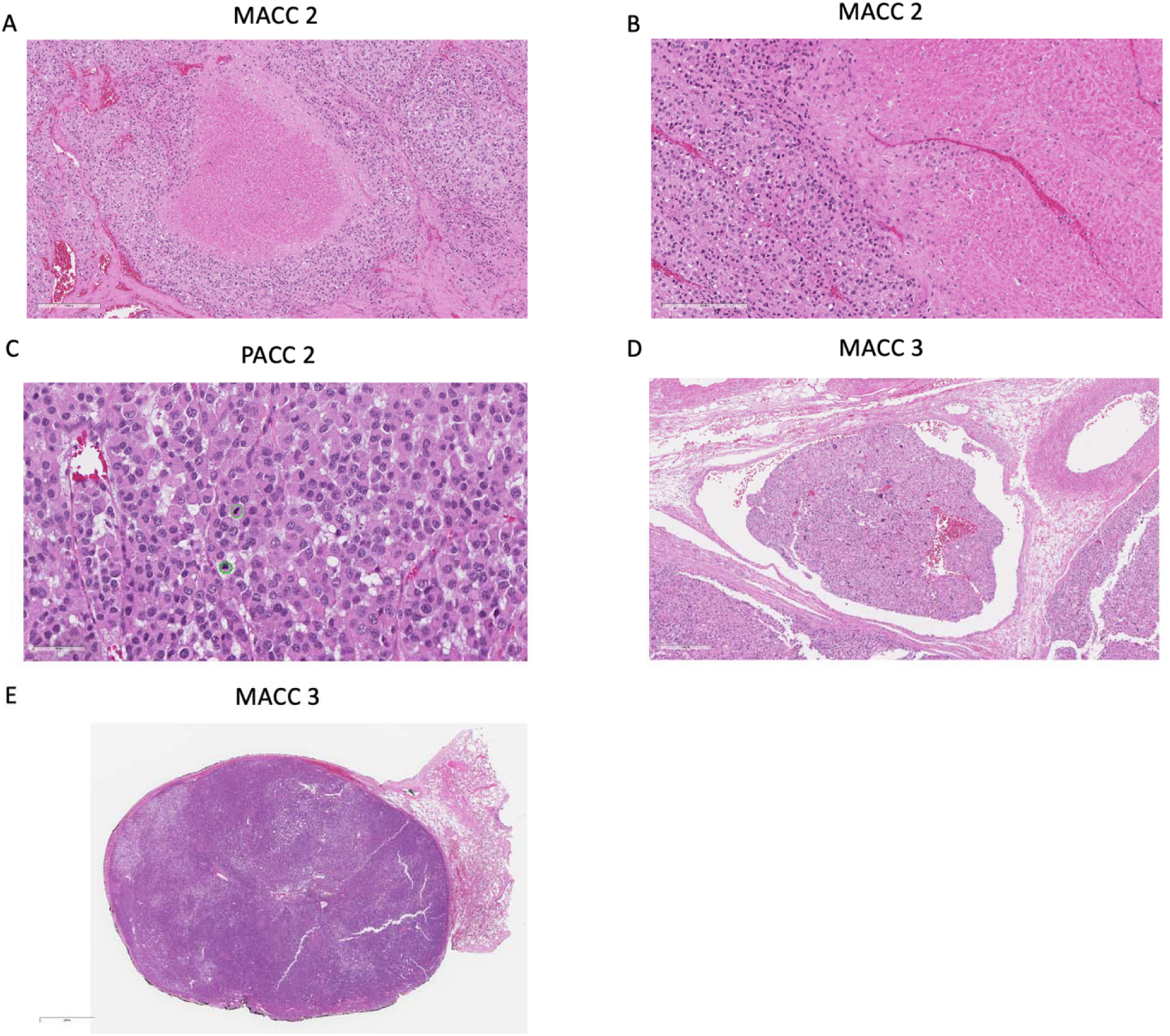
H&E Staining of FFPE Patient Tissues. A. H&E staining of ACC with necrosis (sample MACC 2) under 10X magnification. B. H&E staining of ACC with necrosis (sample MACC 2) under 20X magnification. C. H&E staining of ACC (sample PACC 2) under 40X magnification. Mitoses are highlighted by green circles. D. H&E staining demonstrating ACC with lymphovascular invasion (sample MACC 3) under 10X magnification. E. H&E staining of ACC lung metastasis (sample MACC 3) under 1X magnification.

**Supplementary Figure S8.**
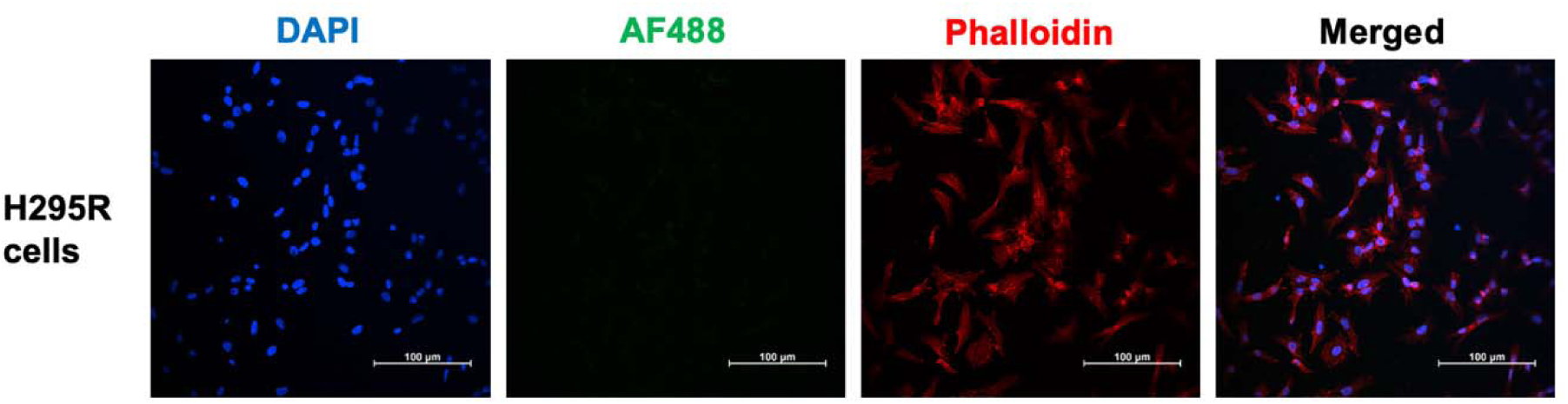
Negative Controls for NCI-H295R Cell Line Immunofluorescence. NCI-H295R cells were processed alongside those shown in Figure 6 but without primary mouse anti-γ-H2AX antibody. The green channel represents background produced by Alexa Fluor® 488-conjugated secondary antibody.

**Supplementary Figure S9.**
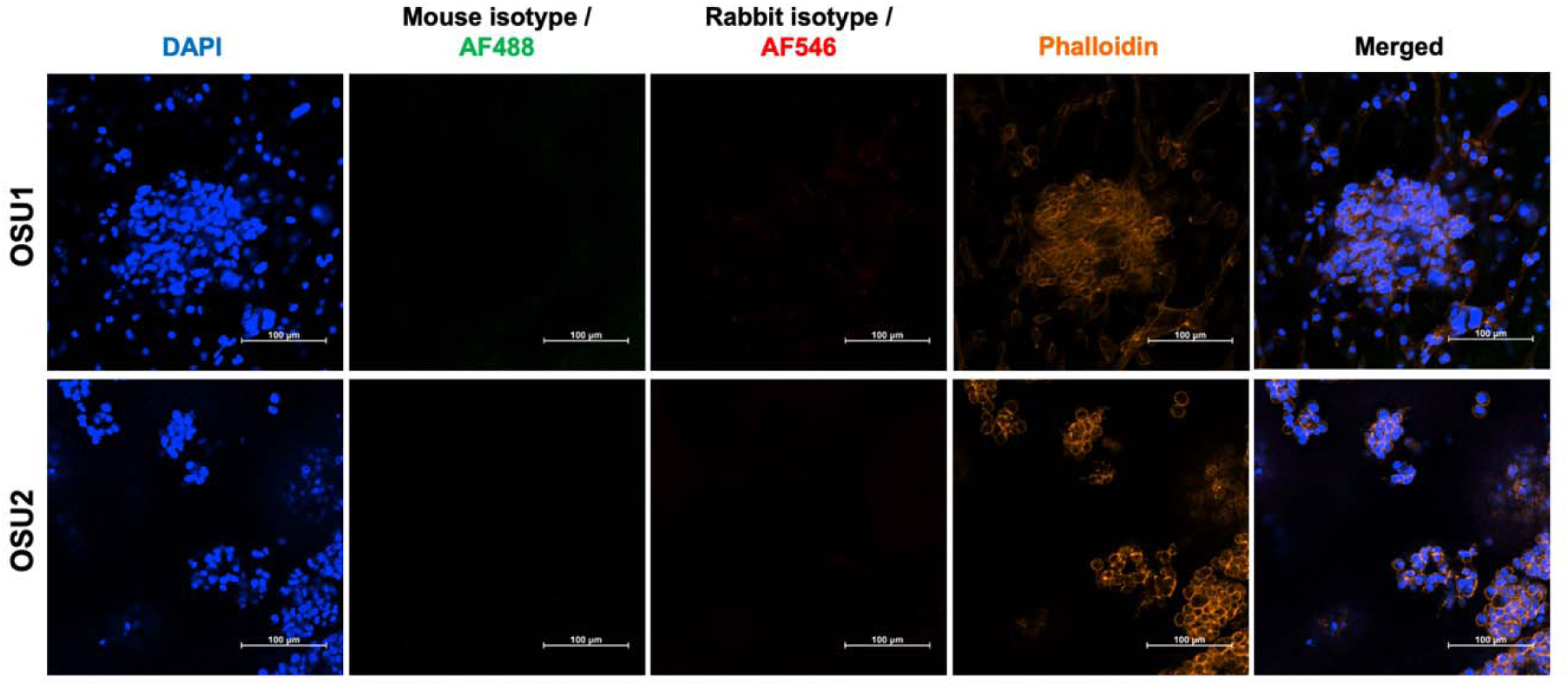
Isotype Controls ACC patient-derived tumor organoid (PTO) Immunofluorescence. ACC PTOs were processed alongside those shown in Figure 7, but instead with anti-rabbit (corresponding to SF-1) and anti-mouse (corresponding to γ-H2AX) primary antibody isotype controls. The red and green channels background staining produced by anti-rabbit Alexa Fluor® 488-and anti-mouse-Alexa Fluor® 546 conjugated secondary antibodies.

**Supplementary Table S1.**
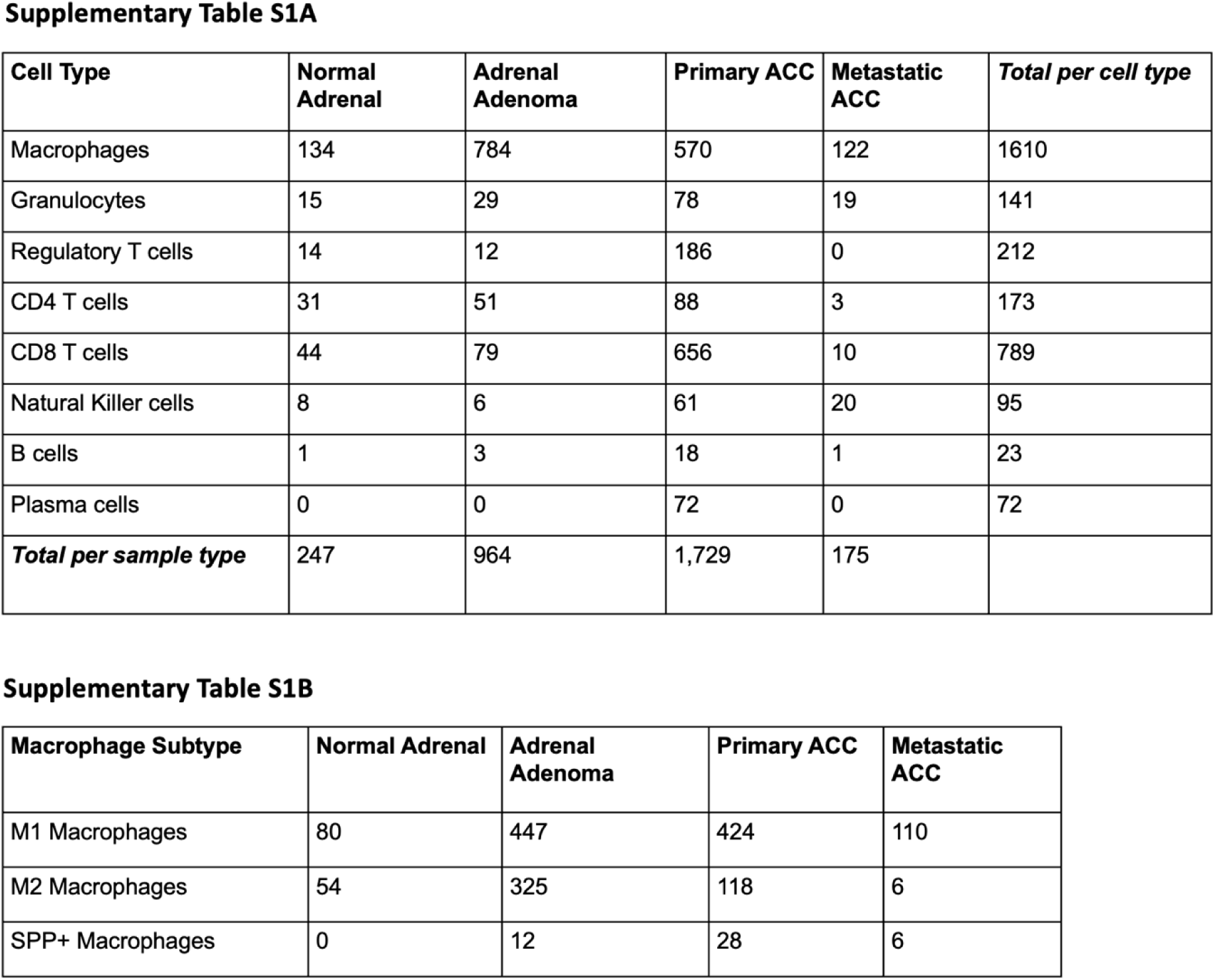
Immunocyte Counts per Sample Type. A. Immunocyte counts per sample type. B. Macrophage subtype counts per sample type.

**Supplementary Table S2.**
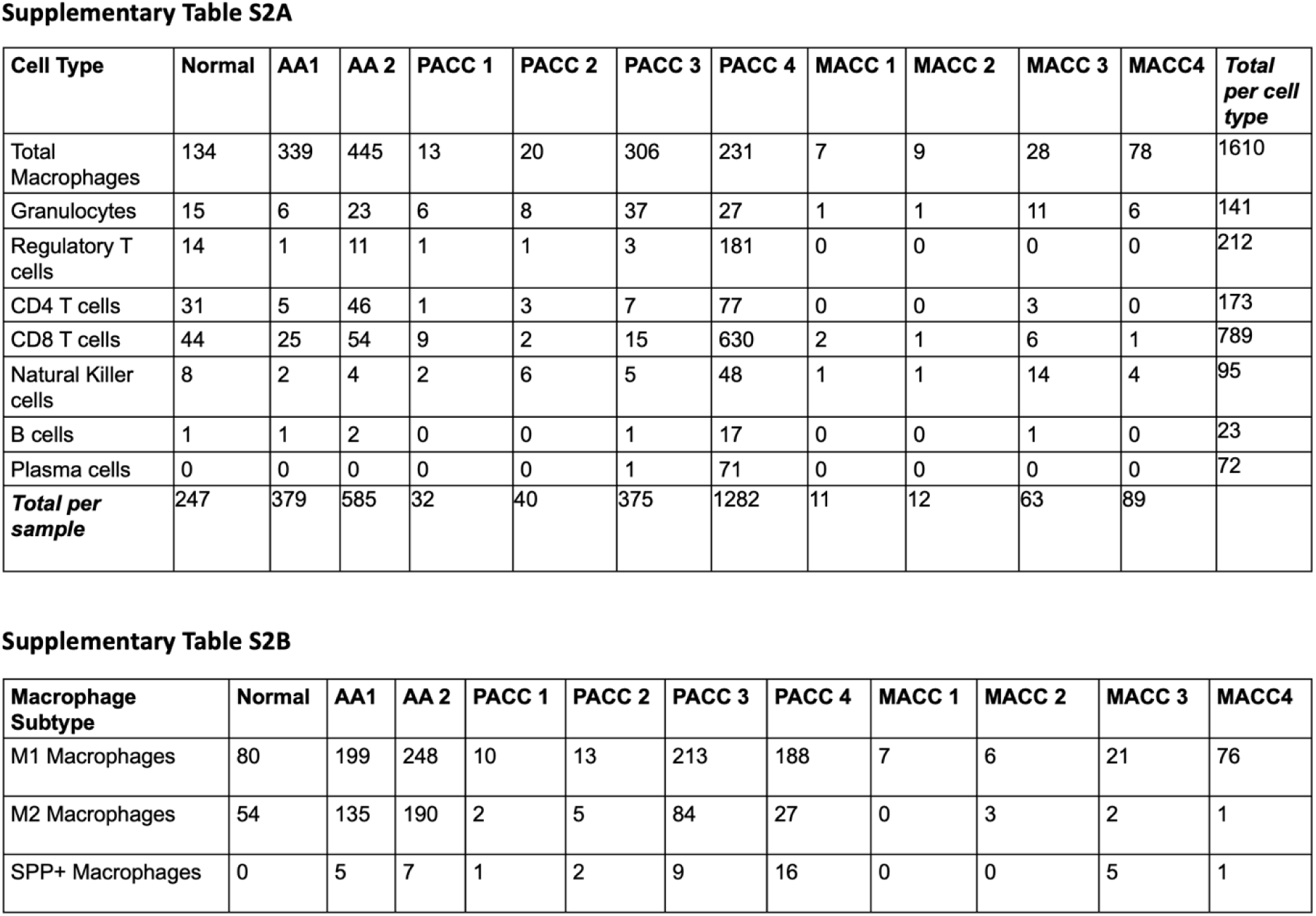
Immunocyte Counts per Sample. A. Immunocyte counts per individual sample. B. Macrophage subtype counts per individual sample.

**Supplementary Table S3.** Summary of Pathology Findings in ACC Specimens. Pathology findings are summarized for samples PACC 2, MACC 2, and MACC 3.

| Samples | Pathology Findings |
| --- | --- |
| PACC 2 | Mitotic count 25/50HPF, Conventional ACC, Capsular, extraglandular and LVI, necrosis 10-20% |
| MACC 2 | Mitotic count 74/ 50 hpf; Oncocytic and myxoid morphology, necrosis (5-10%); capsular and extra-glandular invasion |
| MACC 3 | Mitotic count 25/50HPF, Conventional ACC, Capsula invasion, necrosis 5-10% |

